# CEP signaling coordinates plant immunity with nitrogen status

**DOI:** 10.1101/2022.12.20.521212

**Authors:** Jakub Rzemieniewski, Henriette Leicher, Hyun Kyung Lee, Caroline Broyart, Shahran Nayem, Christian Wiese, Julian Maroschek, Zeynep Camgöz, Vilde Olsson Lalun, Michael Anthony Djordjevic, A. Corina Vlot, Ralph Hückelhoven, Julia Santiago, Martin Stegmann

**Author notes:** Biotechnology of Natural Products, TUM School of Life Sciences, Technical University of Munich, Freising, Germany. Chair of Crop Plant Genetics, Faculty of Life Sciences: Food, Nutrition and Health, University of Bayreuth, Kulmbach, Germany (A. Corina Vlot).

## Abstract

Plant endogenous signaling peptides shape growth, development and adaptations to biotic and abiotic stress. Here, we identified C-TERMINALLY ENCODED PEPTIDEs (CEPs) as novel immune-modulatory peptides (phytocytokines) in *Arabidopsis thaliana*. Our data reveals that CEPs induce immune outputs and are required to mount resistance against the leaf-infecting bacterial pathogen *Pseudomonas syringae* pv*. tomato*. We show that effective immunity requires CEP perception by tissue-specific CEP RECEPTOR 1 (CEPR1) and CEPR2. Moreover, we identified the related RECEPTOR-LIKE KINASE 7 (RLK7) as a novel CEP4-specific CEP receptor contributing to CEP-mediated immunity, suggesting a complex interplay of multiple CEP ligands and receptors in different tissues during biotic stress. CEPs have a known role in the regulation of root growth and systemic nitrogen (N)-demand signaling. We now provide evidence that CEPs and their receptors promote immunity in an N status-dependent manner, suggesting a previously unknown molecular crosstalk between plant nutrition and cell surface immunity. We propose that CEPs and their receptors are central regulators for the adaptation of biotic stress responses to plant-available resources.

## Main Text

Receptor kinases (RKs) sense external and internal cues to control multiple aspects of plant physiology, ranging from growth and development to plant immunity and abiotic stress tolerance. RKs can serve as pattern recognition receptors (PRRs) to detect microbe-associated molecular patterns (MAMPs) and activate pattern-triggered immunity (PTI). An example is the *Arabidopsis thaliana* (hereafter Arabidopsis) leucine-rich repeat RK (LRR-RK) FLAGELLIN SENSITIVE 2 (FLS2), which forms a receptor complex with BRASSINOSTEROID INSENSITIVE 1-ASSOCIATED RK 1 (BAK1) upon perception of a 22 amino acid epitope derived from bacterial flagellin (flg22) to activate PTI (*1–4*). Plants also perceive endogenous peptides to regulate multiple aspects of plant physiology (*5*). Importantly, plants utilize specific endogenous peptides for controlling immunity. These peptides are referred to as phytocytokines and their expression or secretion can be modulated upon PTI activation. They also often additionally regulate aspects of plant growth and development (*6*, *7*). Examples are GOLVEN2 (GLV2) peptides which are perceived by ROOT MERISTEM GROWTH FACTOR INSENSITIVE 3 (RGI3) to modulate PRR stability and RAPID ALKALINIZATION FACTORs (RALFs) that are sensed by the malectin RK (MLRK) FERONIA (FER) to control PRR nanoscale dynamics at the plasma membrane and MAMP-induced PRR-BAK1 complexes for PTI initiation (*8–11*). GLV2 also controls hypocotyl gravicurvature (*12*) and RALF perception by FER and other MLRKs affects several aspects of plant growth, development and reproduction, suggesting that endogenous peptides coordinate these processes with stress responses (*13–18*).

Immune-modulatory peptides are often transcriptionally upregulated in response to MAMP perception, including SERINE-RICH ENDOGENOUS PEPTIDES (SCOOPs) and SMALL PHYTOCYTOKINES REGULATING DEFENSE AND WATER LOSS (SCREWs)/CTNIPs (*19–21*). Yet, *GLV2* transcription is not induced by biotic stress (*8*) and promotes immunity, suggesting that immunity-dependent transcriptional regulation is not a prerequisite for phytocytokine function. Phytocytokines and other endogenous peptides further regulate a multitude of abiotic stress responses, including adaptation to high salinity, drought and nutrient deprivation, indicating that they can integrate multiple external and internal cues to safeguard plant health (*22–24*). Yet, how different peptide-mediated pathways are coordinated remains largely unknown.

Here, we identified C-TERMINALLY ENCODED PEPTIDES (CEPs) as novel phytocytokines in Arabidopsis. CEPs are important for sucrose-dependent lateral root growth, root system architecture, systemic nitrogen (N)-demand signaling and promotion of root nodulation, but a function in plant immunity remained unknown (*22*, *25–31*). We show that the unusual class I CEP peptide CEP4 induces immune responses. We found that *CEP4* and other *CEPs* are expressed in shoots and perceived by canonical CEP receptors CEPR1 and CEPR2 to mount effective cell surface immunity. *CEPR1* and *CEPR2* show tissue-specific expression patterns, suggesting CEP sensing in distinct tissues spatially cooperates to control plant immunity. Yet, CEP4-induced responses also require the CEPR-related RECEPTOR-LIKE KINASE 7 (RLK7), which we identified as a novel CEP4-specific CEP receptor with wide-spread expression in leaves. Importantly, we now show that a short-term reduction in seedling N levels promotes flg22-induced PTI responses in a CEP and CEP receptor-dependent manner, suggesting that CEPs coordinate a previously unknown cross-talk between cell surface immunity and plant nutrition.

## Results

### CEPs are novel phytocytokines

We sought to identify novel phytocytokines regulating growth and immunity in Arabidopsis and screened publicly available transcription data of known growth-regulatory plant peptide families for members with differential expression after elicitor treatment. With this approach, we recently identified GLV2 as a novel phytocytokine modulating PRR stability through RGIs (*8*). We noticed that a specific member of the CEP family, *CEP4*, showed differential expression upon flg22 treatment with a moderate downregulation in an *asr3* mutant background, a transcriptional repressor of flg22-induced genes (Supplementary Fig. 1A) (*32*). Using RT-qPCR, we also observed a mild flg22-induced *CEP4* downregulation in Col-0 seedlings compared to the mock control. At 4 hours of mock treatment, *CEP4* expression was lower compared to 0-hour mock samples, suggesting some degree of circadian rhythm-related regulation (Supplementary Fig. 1B). The majority of *CEPs* show predominant expression in root tissue (*33*). We compared root and shoot *CEP4* transcripts in seedlings upon flg22 treatment. We detected much stronger *CEP4* expression in roots with a similar pattern of flg22-dependent modulation. Interestingly, *CEP4* expression in shoots was mildly upregulated upon 1 h of flg22 treatment (Supplementary Fig. 1C). Arabidopsis encodes for 12 class I CEPs and 3 class II CEPs, which can be distinguished by sequence differences in their peptide domain. CEPs are produced from larger peptide precursors that carry one to five predicted mature CEP domain in their sequence and an N-terminal signal peptide for secretion (*33*). CEP4 is classified as a class I CEP but has an unusual structure compared to other class I CEPs (Supplementary Fig. 1D). CEP4 only carries two proline residues in its peptide domain, unlike several characterized typical class I CEPs such as CEP1 and CEP3 (*33*).

To test whether CEP4 may be involved in immunity, we generated constitutive overexpression lines using a full-length CEP4 precursor sequence (*35S::CEP4*, Supplementary Fig. 2A) and noticed that these lines showed increased resistance to infection by *Pseudomonas syringae* pv. *tomato (Pto)* lacking the effector molecule coronatine (*Pto*^COR-^), which is routinely used to assess PTI-associated disease resistance phenotypes (Fig. 1A) (*34*, *35*). The same lines were also more susceptible to infection by the wild-type *Pto* DC3000 strain (Supplementary Fig. 2B).

**Figure 1:**
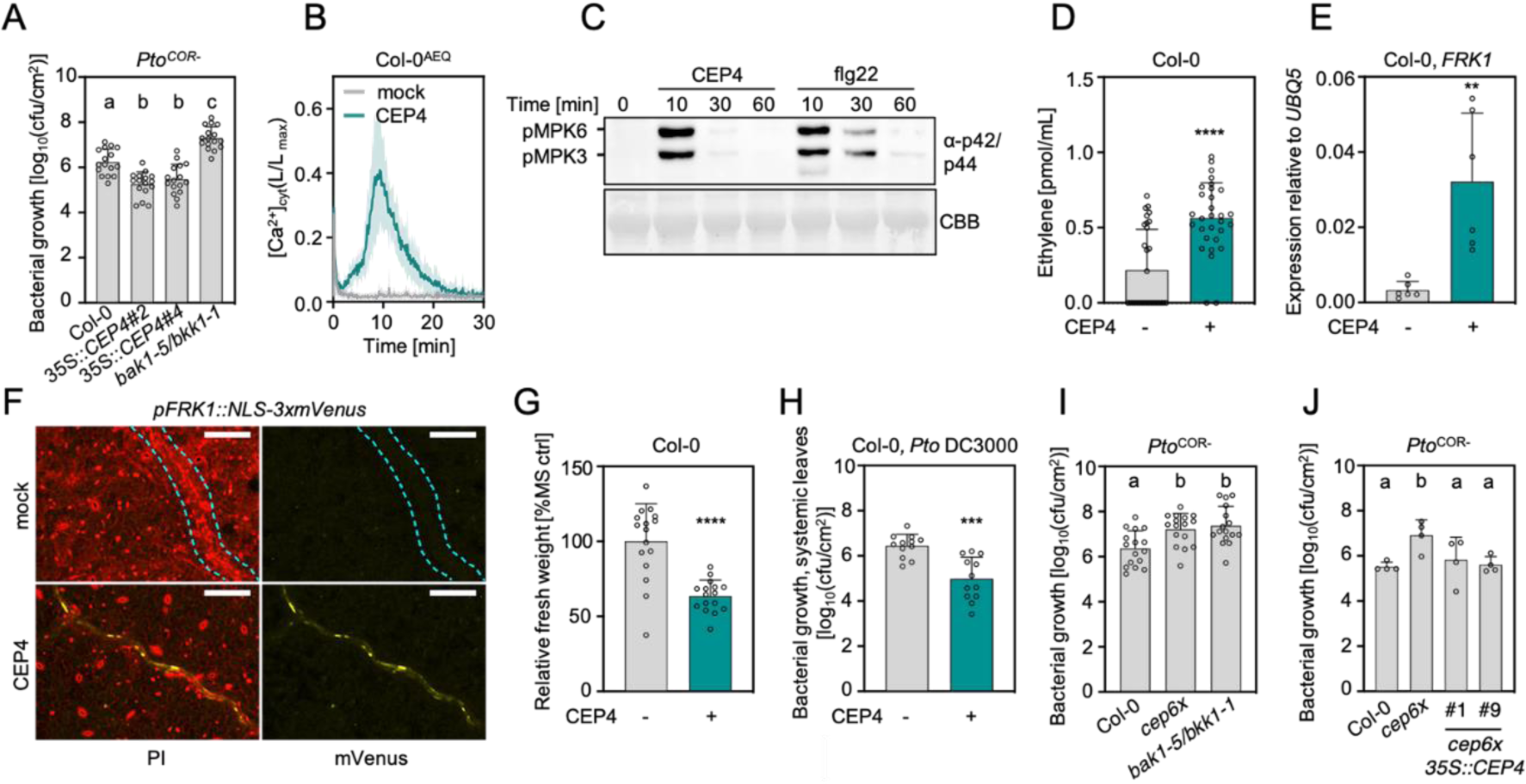
CEPs induce immune responses and are important determinants of plant immunity. **A)** Colony forming units (cfu) of *Pto*^COR-^ 3 days post inoculation (3 dpi) upon spray infection; n = 16 pooled from four experiments ± SD (one-way ANOVA, Tukey post-hoc test; a-b, p<0.01; a/b-c p<0.0001). **B**) Kinetics of cytosolic calcium concentrations ([Ca^2+^]_cyt_) in Col-0^AEQ^ seedlings upon mock (ddH_2_O) or CEP4 (1 μM) treatment; n = 8, ± SD. **C**) MAPK activation in Col-0 upon CEP4 (1 μM) or flg22 (100 nM) treatment for the indicated time. Western blots were probed with α-p44/42. CBB = Coomassie brilliant blue. **D**) Ethylene concentration in Col-0 leaf discs 3.5 h upon mock (ddH_2_O) or CEP4 (1 μM) treatment; n = 30 pooled from six experiments ± SD (two-tailed Student’s t-test, **** p<0.0001). **E**) RT-qPCR of *FRK1* in seedlings upon CEP4 (1 μM) or mock (ddH_2_O) treatment for 4 h. Housekeeping gene *UBQ5*; n = 6 pooled from six experiments ± SD (two-tailed Student’s t-test, ** p<0.01). **F**) NLS-3xmVenus signal in p*FRK1::NLS-3xmVenus* upon mock (ddH_2_O) or CEP4 (100 nM) treatment for 16 h. Cyan dotted line indicates vasculature. PI = propidium iodide, scale bar = 100 μm. **G**) Relative fresh weight (as percent of ½ MS medium control = % MS ctrl) of five-day-old seedlings treated with CEP4 (1 μM) for seven days (n=16, ± SD, **** p<0.0001, two-tailed Student’s t-test). **H**) cfu of *Pto* DC3000 (4 dpi) in distal leaves upon local CEP4 (1 µM) or mock (ddH_2_O) pre-treatment; n = 12 pooled from three experiments ± SD (two-tailed Student’s t-test, *** p=0.0001). **I**) cfu of *Pto*^COR-^ (3 dpi) upon spray infection. The *bak1-5/bkk1-1* mutant was used as a hypersusceptible control; n = 16 pooled from four experiments ± SD (one-way ANOVA, Tukey post-hoc test, a-b p≤0.001). **J**) cfu of *Pto*^COR-^ (3dpi) upon spray infection; n = 4 ± SD (one-way ANOVA, Tukey post-hoc test, a-b p<0.05). All experiments were repeated at least three times in independent biological repeats with similar results.

The majority of mature CEPs previously identified are 15mer peptides with an N-terminal aspartate (D), a C-terminal Histidine (H) and hydroxylated prolines (*22*, *33*). We synthesized a 16mer peptide with both proline residues hydroxylated and the N- and C-terminal D and H residue, respectively, DAFRHypTHQGHypSQGIGH, to test whether it triggered or modulated immune responses. CEP4 application activated dose-dependent PTI outputs, including the cellular influx of calcium ions in a Col-0 line expressing the calcium reporter Aequorin (Col-0^AEQ^) (*36*), the activation of MITOGEN-ACTIVATED PROTEIN KINASEs (MAPKs), ethylene production and expression of the PTI marker gene *FLAGELLIN-INDUCED RECEPTOR KINASE 1* (*FRK1*) in Col-0 seedlings (Fig. 1B-E, Supplementary Fig. 3A). CEP4-induced calcium influx was detectable in the low nanomolar range of CEP4 concentration (Supplementary Fig. 3A) and MAPK phosphorylation was detected at concentrations of 100 nM, yet the magnitude of response was weaker compared to flg22 (Supplementary Fig. 3B). *FRK1* transcript accumulation and calcium influx activated by flg22 treatment was much stronger compared to CEP4 in whole seedlings (Supplementary Fig. 3C, D). However, CEP4 induced *FRK1* expression in a similar range as previously described elicitors, suggesting biological relevance (*37*). Moreover, CEP4 triggered nuclear YFP fluorescence in the vasculature of *pFRK1::NLS-3xmVenus* seedlings, suggesting some degree of tissue specificity of CEP4-induced immune outputs (*38*) (Fig. 1F). Finally, CEP4 treatment resulted in seedling growth inhibition (SGI) and systemic resistance to *Pto* DC3000 infection (Fig. 1G, H). We also tested a 15mer peptide lacking the N-terminal D residue (AFRHypTHQGHypSQGIGH), which triggered a dose-dependent calcium influx with no significant differences to the response elicited by the 16mer peptide (Supplementary Fig. 3A). We next tested whether other class I CEPs can activate PTI responses. Indeed, CEP1 and one peptide derived from CEP9 (CEP9.5), which carries five CEP domains in its precursor sequence (*39*), were able to trigger calcium influx in seedlings but only at higher concentrations of 10 μM (Supplementary Fig. 3E). However, CEP1 induced nuclear YFP fluorescence in the vasculature of *pFRK1::NLS-3xmVenus* lines at similar concentrations as CEP4 (Supplementary Fig. 3F), suggesting that likely several CEPs can trigger immune responses with CEP4 being a very potent family member.

To confirm the role of CEPs in plant immunity and because we anticipated genetic redundancy of immune-regulatory CEPs, we generated a *cep6x* mutant by CRISPR-Cas9 in which *CEP4*, as well as the five additional class I CEPs *CEP1*-*CEP3*, *CEP6* and *CEP9* were mutated to predictable loss of function (CRISPR alleles *cep1.1*, *cep2.1*, *cep3.1*, *cep4.1*, *cep6.1* and *cep9.1*, Supplementary Fig. 2C). The resulting *cep6x* mutant had no obvious morphological defects (Supplementary Fig. 2D) but showed compromised resistance to *Pto^COR-^* infection, confirming that CEPs are important for antibacterial resistance (Fig. 1I). To overcome the impact of different tissue-specific CEPs that cooperate to mount disease resistance, we complemented the *Pto*^COR-^ hypersusceptibility phenotype of *cep6x* using a full-length *CEP4* driven by the constitutive 35S promoter (Fig. 1J, Supplementary Fig. 2E). The *cep6x* mutant was also more susceptible to infection with the fully virulent wild-type *Pto* DC3000 strain, and *35S::CEP4* expression partially complemented this phenotype (Supplementary Fig. 4A). We also tested a *cep5x* mutant (CRISPR alleles *cep1.2*, *cep2.2*, *cep3.2*, *cep6.2* and *cep9.2*, *CEP4* wild-type) which showed an intermediate phenotype upon *Pto*^COR-^ infection (Fig. 4B, Supplementary Fig. 2C, D). This indicates the contribution of multiple CEPs to mount robust resistance.

### CEPR1 and CEPR2 are CEP4 receptors and central regulators of plant immunity

CEPs bind to CEPR1 and CEPR2, two LRR-RLKs from LRR subfamily XI (*22*, *40*). Genetically, CEPR1 is predominantly required for class I CEP perception during root growth-related responses (*25–28*, *41*, *42*). We tested whether *CEPR1* and *CEPR2* are also involved in bacterial immunity. We did not observe a *Pto^COR-^* infection phenotype in *cepr1-3* and *cepr2-4* single mutants (Supplementary Fig. 5A, Fig. 2A). We generated a *cepr1-3 cepr2-4 (cepr1-3/2-4*) mutant by genetic crossing and this double mutant was more susceptible to *Pto*^COR-^ (Fig. 2A). We also generated a new *cepr1 cepr2* double mutant by CRISPR-Cas9 in a Col-0^AEQ^ background (CRISPR alleles *cepr1.1*^AEQ^, *cepr2.1*^AEQ^, hereafter *cepr1/2*^AEQ^) (Supplementary Fig. 5B). Similar to *cepr1-3/2-4*, *cepr1/2*^AEQ^ was more susceptible to *Pto*^COR-^ infection (Fig. 2A), further confirming a role of *CEPR1* and *CEPR2* in antibacterial resistance. These data suggest that *CEPR1* and *CEPR2* may control immunity redundantly.

**Figure 2:**
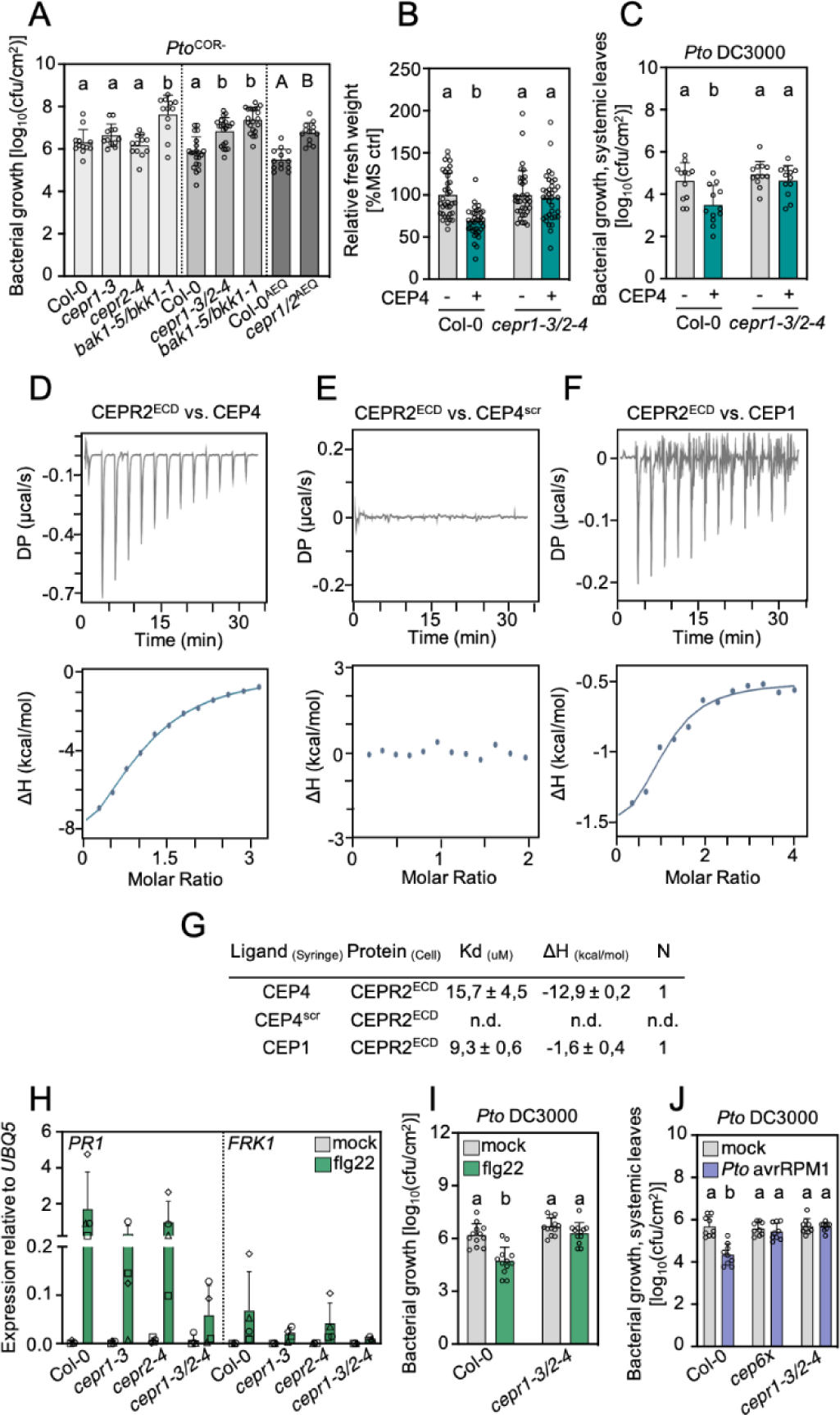
CEPR1 and CEPR2 are CEP4 receptors and are important determinants of plant immunity. **A)** cfu of *Pto*^COR-^ (3 dpi) upon spray infection. The dotted line indicates different experiments; n = 12, 21 and 13 pooled from three, five and three experiments, respectively ± SD (one-way ANOVA, Dunnett’s post-hoc test, a-b p<0.0001, two-tailed Student’s t-test, A-B p<0.0001). **B)** Relative fresh weight of five-days-old seedlings treated with CEP4 (1 μM) for seven days; n = 35-36 pooled from three experiments ± SD (one-way ANOVA, Tukey post-hoc test, a-b p<0.0001). **C)** cfu of *Pto* DC3000 (4 dpi) in distal leaves upon local CEP4 (1 µM) or mock (ddH_2_O) pre-treatment; n = 12 pooled from three experiments ± SD (one-way ANOVA, Tukey post-hoc test, a-b p<0.01). **D)** CEP4, **E**) CEP4^scr^ and **F**) CEP1 were titrated into a solution containing CEPR2^ECD^ in ITC cells. Top: raw data thermogram; bottom: fitted integrated ITC data curves. DP = differential power between reference and sample cell; ΔH = enthalpy change. **G)** ITC table summarizing CEPR2^ECD^ vs CEP4/CEP4^scr^/CEP1 as means ± SD of two experiments. The dissociation constant (K_d_) indicates receptor-ligand binding affinity. N indicates reaction stoichiometry (n = 1 for 1:1 interaction). **H)** RT-qPCR of *PR1* and *FRK1* in adult leaves after treatment with flg22 (1 μM) or mock (ddH_2_O) for 24 h. Housekeeping gene *UBQ5*. Different symbols represent four independent experiments; n = 4 ± SD. **I)** cfu of *Pto* DC3000 (3 dpi) in leaves upon flg22 (1 µM) or ddH_2_O pre-treatment; n = 12 pooled from three experiments ± SD (one-way ANOVA, Tukey post-hoc test, a-b p<0.0001). **J)** cfu of *Pto* DC3000 (4 dpi) in distal leaves upon local infection with *Pto avrRPM1* or 10 mM MgCl_2_; n = 9 pooled from three independent experiments ± SD (one-way ANOVA, Tukey post-hoc test, a-b p<0.0001). Experiments in **A**-**C**, **H-J,** and **D**-**F** were repeated at least three times in independent biological repeats or in two independent technical repeats, respectively, with similar results.

We then tested whether CEP4 perception depends on CEPR1 and/or CEPR2. Using SGI as a readout, we noticed that *cepr1-3* was insensitive and two *cepr2* alleles (*cepr2-3* and *cepr2-4*) were insensitive and less sensitive to CEP4 treatment, respectively (Supplementary Fig. 5C). The *cepr1-3/2-4* mutant and the previously reported No-0 *cepr1-1 cepr2-1* double mutant also did not respond to CEP4 in SGI experiments (Fig. 2B, Supplementary Fig. 5D). Similarly, the *cepr1-3/2-4* double mutant was largely insensitive to CEP4-induced systemic resistance (Fig. 2C). Of note, mock-treated *cepr1-3/2-4* mutants do not show enhanced bacterial growth compared to Col-0, suggesting that CEPR1/CEPR2 primarily regulate immunity to tissue invasion, an effect that is bypassed by syringe infiltration in this experimental setup. Collectively, these data suggest that both CEPR1 and CEPR2 are involved in CEP4 perception. We then tested whether CEP4 can directly bind to the ectodomain (ECD) of CEPR1 and/or CEPR2. We expressed CEPR1^ECD^ and CEPR2^ECD^ in *Trichoplusia ni* Tnao38 cells and purified them for quantitative binding experiments. We analyzed protein quality by Coomassie stain and size exclusion chromatography (SEC, Supplementary Fig. 6A, B). Unfortunately, CEPR1^ECD^ aggregated, as indicated by the early elution of the bulk sample during SEC analysis (∼10min, Supplementary Fig. 6A). Nevertheless, we obtained good quality protein for CEPR2^ECD^ with a single SEC elution peak at ∼13min, which we subsequently tested for quantitative binding to CEP4 using isothermal titration calorimetry (ITC). CEP4, but not a scrambled control (CEP4^scr^), directly bound to CEPR2^ECD^ with a K_D_ of 15.7 μM (± 4.5 μM) (Fig. 2D-E, G, Supplementary Fig. 6C). CEP1, which was previously shown to bind to CEPR1 and CEPR2 (*22*) also bound to CEPR2^ECD^ with a K_D_ of 9.3 μM (± 0.6 μM) (Fig. 2F-G, Supplementary Fig. 6C). These data are in range with previously reported peptide-LRR-RK binding affinities obtained by ITC and suggest that CEPR2 is a bona fide CEP4 receptor (*43–45*).

We were then interested to characterize a possible role of CEPR1 and CEPR2 for FLS2-mediated signaling. We did not observe strong differences in flg22-induced ethylene accumulation *in cepr1-3/2-4* (Supplementary Fig. 7A). Of note, flg22-induced ethylene production was higher in *cepr1-3/2-4*, but basal ethylene production was enhanced in this mutant background, making the result difficult to interpret. This is interesting in light of a previous report showing that the *Medicago truncatula* CEP1-CRA2 (CEPR1 orthologue) pathway negatively regulates ethylene signaling during root nodule symbiosis (*46*). Interestingly though, flg22-induced *FRK1* and *PR1* expression were reduced in adult *cepr1-3* or *cepr2-4* plants, which was pronounced in *cepr1-3/2-4* (Fig. 2H). Moreover, *cepr1-3/2-4* showed compromised flg22-induced resistance to *Pto* DC3000 infection (Fig. 2I). The flg22-induced seedling growth inhibition was unaffected in *cepr1-3/2-4* (Supplementary Fig. 7B), suggesting that CEPR1 and CEPR2 are selectively required for specific flg22-induced outputs associated with antibacterial defense. We next tested whether CEP-CEPR1/2 signaling may be important for systemic acquired resistance (SAR). To induce SAR, we used a *Pto* strain producing the effector avrRPM1 (*Pto* avrRPM1), which is recognized by Col-0 RESISTANCE TO PSEUDOMONAS SYRINGAE 1, to activate effector-triggered immunity and consequently SAR (*47*, *48*). Local inoculation of *Pto avrRPM1* and subsequent infection of systemic tissue with virulent *Pto* revealed that *cepr1-3/2-4*, as well as *cep6x*, were strongly compromised in *Pto* avrRPM1-triggered SAR (Fig. 2J). Collectively, these data show that CEP-CEPR signaling is a central regulator of PTI and SAR in Arabidopsis.

### RLK7 is a novel CEP4-specific CEP receptor

To further characterize the role of CEPR1 and CEPR2 in CEP4-induced signaling, we tested early CEP4-triggered responses in *cepr1-3/2-4* and *cepr1/2*^AEQ^. To our surprise, we found that *cepr1/2*^AEQ^ did not show compromised calcium influx upon CEP4 treatment (Fig. 3A). CEP4-induced MAPK activation was also unaffected in *cepr1-3/2-4* (Fig. 3B). Moreover, *cepr1-3* and *cepr2-4* were largely unaffected in CEP4-induced *FRK1* expression and the response was also not abolished in *cepr1-3/2-4* (Supplementary Fig. 8A). These results suggested that other receptor(s) may be involved in CEP4 perception. CEPR1 and CEPR2 are phylogenetically close to IKU2, which is involved in seed size regulation, and RLK7, which plays a role in controlling germination speed, lateral root formation and salt stress adaptation (*24*, *40*, *49–51*). RLK7 also senses endogenous PAMP-INDUCED PEPTIDES (PIPs) to regulate PTI and resistance to the fungal wilt pathogen *Fusarium oxysporum* and *Pto* (*52*, *53*). We tested whether *iku2-4*, *rlk7-1* and *rlk7-3* are compromised for CEP4 perception. The *iku2-4* mutant showed unaltered CEP4-induced ethylene accumulation, but this response was compromised in *rlk7-1* and *rlk7-3* (Supplementary Fig. 8B). Similarly, *rlk7-1* and *rlk7-3* showed strongly reduced CEP4-induced MAPK activation (Fig. 3C). We also generated an *rlk7/iku2*^AEQ^ mutant by CRISPR-Cas9 in a Col-0^AEQ^ background (CRISPR alleles *rlk7.1*^AEQ^ and *iku2.1*^AEQ^, Supplementary Fig. 8C). The *rlk7/iku2*^AEQ^ line showed compromised CEP4-induced calcium influx (Supplementary Fig. 8D). Two additional *rlk7*^AEQ^ single mutants generated by CRISPR-Cas9 (CRISPR alleles *rlk7.2*^AEQ^*, rlk7.3*^AEQ^

**Figure 3:**
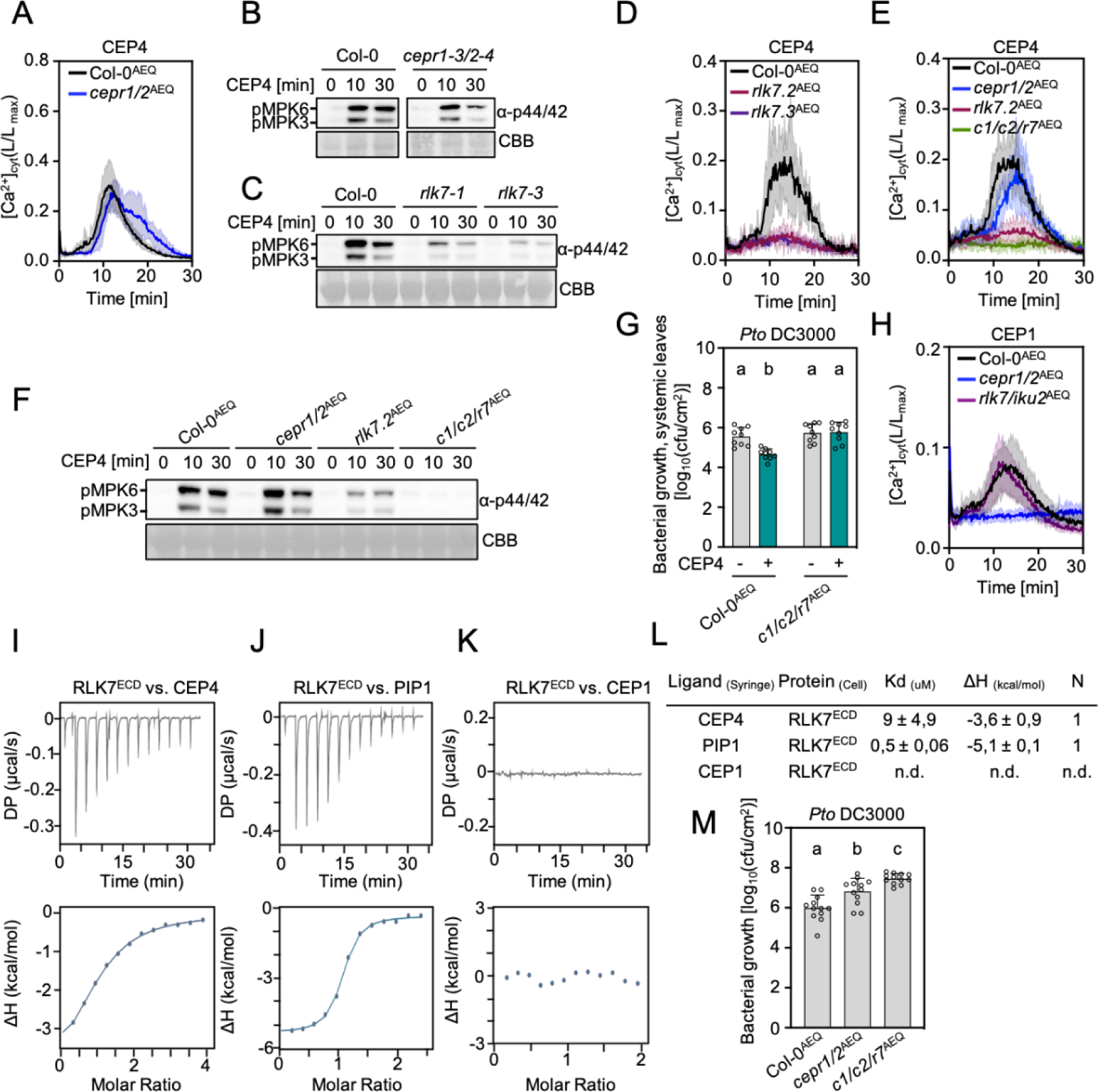
RLK7 is an additional CEP4 receptor. **A)** ([Ca^2+^]_cyt_) kinetics in seedlings upon CEP4 (1 μM) treatment; n = 6 ± SD. **B**-**C)** MAPK activation upon CEP4 (1 μM) treatment for the indicated time. Western blots were probed with α-p44/42. CBB = Coomassie brilliant blue. **D**-**E)** [Ca^2+^]_cyt_ kinetics in seedlings upon CEP4 treatment (1 μM); n = 3, n = 4, respectively ± SD. *c1c2r7* = *cepr1/2/rlk*7^AEQ^**F)** MAPK activation upon CEP4 (1 μM) treatment for the indicated time. Western blots were probed with α-p44/42. CBB = Coomassie brilliant blue. **G)** cfu of *Pto* DC3000 (4 dpi) in distal leaves upon local CEP4 (5 µM) or mock (ddH_2_O) pre-treatment; n = 9 pooled from three experiments ± SD (one-way ANOVA, Tukey post-hoc test, a-b p<0.01). **H)** [Ca^2+^]_cyt_ kinetics in seedlings upon CEP1 (10 μM) treatment; n = 4, ± SD. **I)** CEP4, **J)** PIP1 and **K)** CEP1 were titrated into a solution containing CEPR2^ECD^ in ITC cells. Top: raw data thermogram; bottom: fitted integrated ITC data curves. DP = differential power between reference and sample cell; ΔH = enthalpy change. **L)** ITC table summarizing RLK7^ECD^ vs CEP4/PIP1/CEP1 as mean ± SD of two experiments. The dissociation constant (K_d_) indicates receptor-ligand binding affinity. N indicates reaction stoichiometry (n = 1 for 1:1 interaction). **M)** cfu of *Pto* DC3000 (3 dpi) upon spray infection; n = 12 pooled from three independent experiments ± SD (one-way ANOVA, Tukey post-hoc test, a-b p<0.0021; a-c p<0.0001; b-c p<0.0209). Experiments in **A**-**H**, **M** and **I**-**K** were repeated at least two times in independent biological repeats or in two independent technical repeats, respectively, with similar results.

Supplementary Fig. 8C) were also compromised in CEP4-triggered calcium influx (Fig. 3D). Yet, residual CEP4 activity remained in *rlk7* mutants, both for CEP4-induced MAPK activation and calcium influx (Fig. 3C, D). To resolve this, we generated a CRISPR *cepr1 cepr2 rlk7*^AEQ^ (hereafter *cepr1/2/rlk7*^AEQ^) triple mutant using the *cepr1/2*^AEQ^ background (CRISPR allele *rlk7.4*^AEQ^ in *cepr1/2*^AEQ^, Supplementary Fig. 5B, 8C), which showed abolishment of CEP4-induced calcium influx and MAPK activation (Fig. 3E, F). This suggests that RLK7, CEPR1 and CEPR2 each participate in mounting a full CEP4 response, with RLK7 playing a predominant genetic role. The *cepr1/2/rlk7*^AEQ^ mutant also showed abolished CEP4-induced resistance to *Pto* DC3000 infection (Fig. 3G). Both *rlk7*^AEQ^ and *cepr1/2/rlk7*^AEQ^ were also insensitive to PIP1 in MAPK activation (Supplementary Fig. 8E), consistent with RLK7’s function as a PIP receptor (*52*). Interestingly, *rlk7-1* and *rlk7-3* were only moderately affected in CEP4-induced SGI (Supplementary Fig. 8F), suggesting that certain CEP4 responses require selective specificity for one of the three CEP receptors. Moreover, CEP1 and CEP9.5-induced calcium influx was abolished in *cepr1/2*^AEQ^ and unaltered *in rlk7/iku2*^AEQ^ or *rlk7.2*^AEQ^, respectively, indicating that CEPR1/2 are required for the early responses triggered by these canonical class I CEPs, which again show differential receptor requirements (Fig. 3H, Supplementary Fig. 8G).

We next used ITC to test whether CEP4 can bind to the ectodomain of RLK7 (RLK7^ECD^). Similar to CEPR2^ECD^, we obtained good quality proteins for RLK7^ECD^ eluting with a main single peak at ∼13min during SEC analysis (Supplementary Fig. 6A, B). CEP4 bound to RLK7^ECD^ with a K_D_ of 9 μM (± 4.9 μM) (Fig. 3I, L, Supplementary Fig. 6C), similar to CEPR2^ECD^ (Fig. 2D, G). We also tested binding of PIP1 to RLK7^ECD^, a described RLK7 ligand (*24*, *52*). RLK7^ECD^ bound PIP1 with a higher affinity (K_D_ 500 nM ± 60 nM) (Fig. 3J, L, Supplementary Fig. 6C). Importantly, consistent with unaltered CEP1-induced responses in *rlk7/iku2^AEQ^* (Fig. 3H), CEP1 did not bind to RLK7 (Fig. 3K, Supplementary Fig. 6C). These data suggest that RLK7 can function as a CEP4-specific CEP receptor.

The *cepr1/2*^AEQ^ mutants show compromised resistance to *Pto*^COR-^ (Fig. 2A), similar to previously published *rlk7* single mutants (*53*). Further mutation of *rlk7* in *cepr1/2*^AEQ^ background did not significantly enhance this phenotype (Supplementary Fig. 8H). When using the fully virulent *Pto* DC3000 strain, *cepr1/2*^AEQ^ was moderately more susceptible (Fig. 3M), whereas *rlk7* was not (*53*). Interestingly, the *cepr1/2/rlk7*^AEQ^ triple mutant showed significantly increased susceptibility to *Pto* DC3000 compared to *cepr1/2*^AEQ^ (Fig. 3M), indicating that all three CEP receptors mount full antibacterial resistance.

### CEP-CEP receptor signaling promotes local immunity against *Pto*

Class I *CEPs* show predominant expression in root tissue (*33*) (Supplementary Fig. 1C) and function as systemic root-to-shoot transmitters of N starvation (*22*). We hypothesized that root-expressed *CEPs* may regulate leaf immunity upon infection with *Pto*. Therefore, we performed reciprocal grafts between Col-0 and *cep6x* and tested whether root or shoot expression of *CEPs* is required for immune regulation. Surprisingly, we found that *CEP* mutation in the shoot conferred increased *Pto^COR-^* susceptibility in *cep6x* (Fig. 4A). We used the hypovirulent *Pto*^COR-^ strain in this experiment, as the *cep6x* phenotype is stronger with this bacterial strain (Fig. 1I, Supplementary Fig. 4A). Consistently, we detected weak *CEP1-CEP4*, *CEP6* and *CEP9* expression in leaf tissue with *CEP4* being slightly upregulated upon *Pto* DC3000 infection (Fig. 4B), similar to the mild *CEP4* upregulation upon flg22 treatment in seedling shoots (Supplementary Fig. 1C). We similarly performed reciprocal grafting with Col-0^AEQ^ and *cepr1/2/rlk7*^AEQ^ mutants, which demonstrated that shoot expressed CEP receptors are required to confer enhanced *Pto* DC3000 resistance (Fig. 4C). *Pto* DC3000 was used because the phenotype of *cepr1/2/rlk7*^AEQ^ was more pronounced with this bacterial strain (Fig. 3M). These data suggest that CEP function in the shoot is necessary for their immune-modulatory function, unlike the root-to-shoot CEP mobility required for N-demand signaling (*22*).

**Figure 4:**
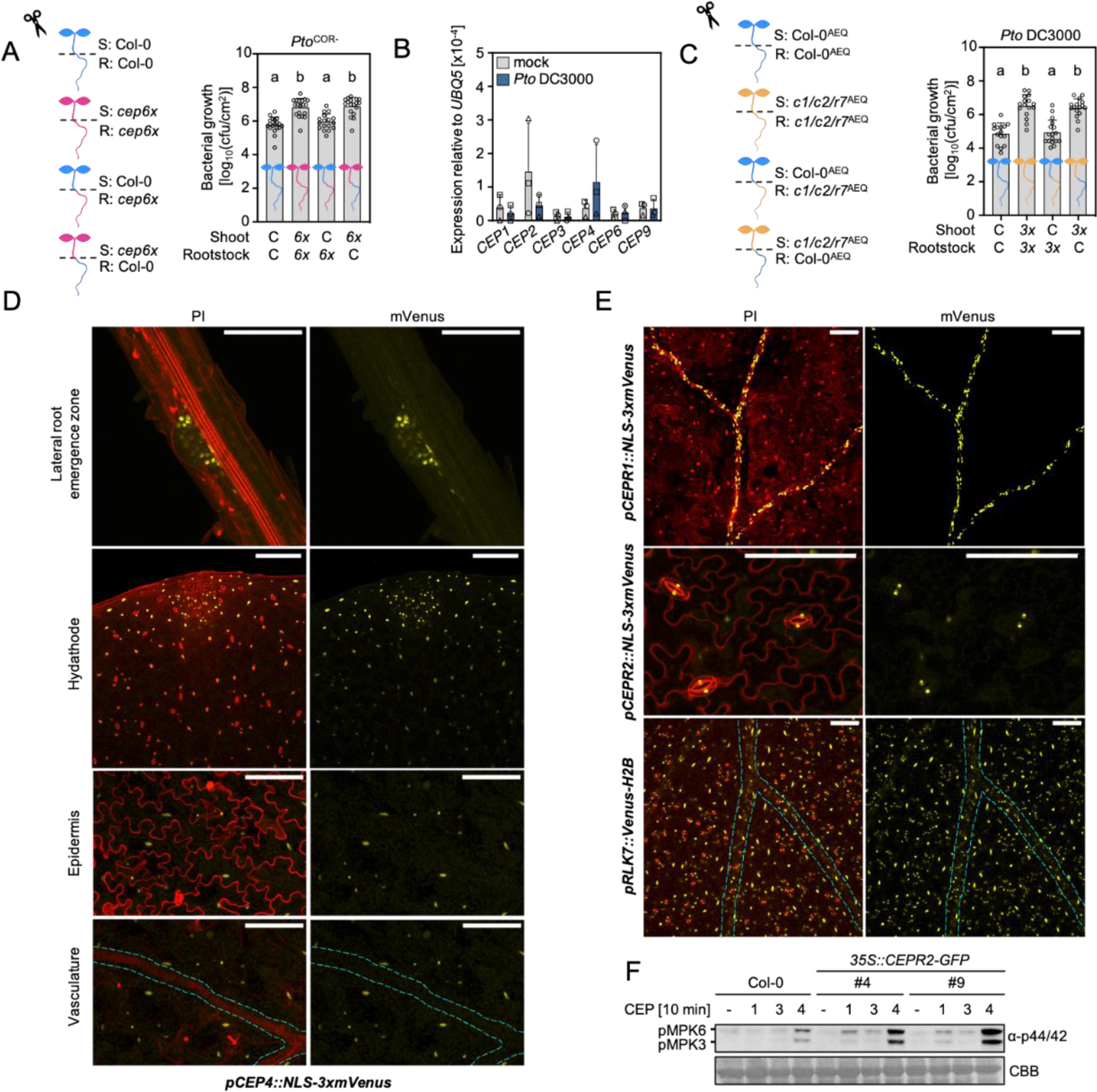
Shoot-expressed *CEPs* and *CEPR1*, *CEPR2*, *RLK7* receptors are required for basal immunity against *Pto*. **A)** Cfu of *Pto*^COR-^ (3 dpi) upon spray infection of reciprocally grafted Col-0 (C) and *cep6x* (*6x*) plants. The genotype of the shoot scion is indicated with S, and the genotype of the rootstock is indicated with R; n = 17-19 pooled from four experiments ± SD (one-way ANOVA, Tukey post-hoc test, a-b p<0.0001). **B)** RT-qPCR analysis of *CEP* expression in mock (ddH_2_O) and *Pto* DC3000-inoculated leaves 24 h post-treatment. Housekeeping gene *UBQ5*; n = 3, ± SD. **C)** cfu of *Pto*^COR-^ (3 dpi) upon spray infection of reciprocally grafted Col-0^AEQ^ (C) and *cepr1/2/rlk7*^AEQ^ (*3x*) plants. The genotype of the shoot scion is indicated with S, and the genotype of the rootstock is indicated with R; n = 15-17 pooled from three experiments ± SD (one-way ANOVA, Tukey post-hoc test, a-b p<0.0001). **D)** pCEP4::NLS-3xmVenus signal in the indicated plant tissues. For imaging the lateral root emergence zone and the hydathode region, the maximum projection of Z-stacks for mVenus is merged with Z-stacked propidium iodide (PI) signal. To show the lack of mVenus signal in the vasculature, the maximum projection of Z-stacks for mVenus is merged with the same single section of PI, showing a single epidermal layer or vasculature (cyan-dotted line). **E)** NLS-3xmVenus or Venus-H2B signal in the leaves of indicated lines. The maximum projection of Z-stacks for mVenus is merged with Z-stacked PI signal. The cyan-dotted line represents vasculature; scale bar = 100 μm. **F)** MAPK activation upon CEP1 (1), CEP3 (3) and CEP4 (4) 1 μM treatment. Western blots were probed with α-p44/42. CBB = Coomassie brilliant blue. Different symbols in **B** represent independent experiments. Experiments were repeated at least two times in independent biological repeats with similar results, except **F**, which has been repeated once with identical results.

Next we wanted to investigate spatial expression patterns of *CEP4* by generating a *pCEP4::NLS-3xmVenus* line. Consistent with previous reports, *CEP4* did not show expression in the main root, but in emerging lateral roots (Fig. 4D) (*33*). Despite weak shoot signals for *CEP4* expression obtained by qPCR (Supplementary Fig. 1C), we found widespread expression of *CEP4* in seedling leaf tissue, but not in the vasculature or stomatal guard cells (Fig. 4D). This data further supports a role for leaf-expressed CEP4 in local responses.

We were next interested in resolving the spatial expression pattern of CEPR1, CEPR2 and RLK7 in shoot tissue. Previous reports using promoter::β-GLUCURONIDASE lines suggested restricted *CEPR1* expression to the vasculature and more widespread *CEPR2* promoter activity (*22*, *54*). Consistently, a *pCEPR1::NLS-3xmVenus* revealed that *CEPR1* expression was specific to vasculature tissue, while *pCEPR2::NLS-3xmVenus* signals were largely restricted to stomatal guard cells (Fig. 4E). The *pRLK7::Venus-H2B* line showed widespread promoter activity, including the vasculature, mesophyll and epidermal cells (Fig. 4E). *RLK7* showed a large overlap with *CEP4* expression in leaves, suggesting that CEP4 and RLK7 can meet *in vivo* to function as a receptor-ligand pair. Although our results point to CEPR1/CEPR2 and RLK7 being required for full CEP4 sensitivity (Fig. 3E, F), only *RLK7* shows an overlapping expression pattern with *CEP4*. Moreover, *CEP4* expression was absent in the vasculature (Fig. 4D) but CEP4 induced *FRK1* expression in this tissue (Fig. 1F, Supplementary Fig. 3F). This raises the possibility for CEP4 mobility between tissue layers and that the peptide may exert its function in a combination of cell-autonomous and short-to-long distance signaling.

The restriction of expression of *CEPR1* and *CEPR2* in the vasculature and guard cells, respectively, may also explain the minor contribution of these receptors to mount CEP4-induced early responses upon elicitation in whole seedlings (Fig. 3A-F). For this reason, we generated *CEPR2-GFP* and *CEPR1-GFP* overexpression lines to test whether the constitutive expression of these receptors can promote CEP4-induced responses. Two *35S::CEPR2-GFP* lines overexpressed the receptor ∼100-150 fold compared to Col-0 and protein accumulation was detected by western blots (Supplementary Fig. 9A, B). We only obtained one *35S::CEPR1-GFP* line with a ∼10 fold overexpression relative to Col-0 (Supplementary Fig. 9C). However, we failed to detect CEPR1-GFP accumulation in this line, suggesting that the protein may be unstable. Consistent with a function as a CEP4 receptor (Fig. 2D, G), the overexpression of CEPR2-GFP enhanced the responsiveness of seedlings to CEP4 in MAPK activation experiments (Fig. 4F). As expected from the lack of detectable protein, *CEPR1* transcript overexpression did not alter CEP4-induced MAPK activation (Supplementary Figure 9D). However, *CEPR1* overexpression mildly promoted CEP3-induced MAPK activation (Supplementary Fig. 9D), in line with previous reports of CEPR1 being the primary receptor for canonical class I CEPs (*22*, *27*, *30*, *31*, *42*). This suggests that CEPR2 is a physiologically relevant CEP4 receptor.

### CEP signaling promotes FLS2 signaling under reduced nitrogen supply

Unlike other immune-promoting phytocytokines, such as PIPs and SCOOPs, *CEP4* is transcriptionally downregulated upon flg22 perception in whole seedling and roots (Supplementary Fig. 1A-C), and only mildly upregulated upon flg22 treatment in seedling shoots or upon *Pto* infection in leaf tissue (Supplementary Fig. 1C, Fig. 4B). We were thus interested to determine the biological relevance of CEP-mediated control of cell surface immunity. First, we tested whether CEP treatment affects FLS2-dependent signaling. CEP4 application could promote flg22-induced ethylene accumulation and resistance induced by a low dose of flg22 (100 nM) (Fig. 5A, B). Yet, we did not observe a noticeable defect in flg22-induced ethylene accumulation in *cep6x* and *cepr1/2/rlk7*^AEQ^ (Supplementary Fig. 10A, B). CEP4 did not induce ethylene production in *cepr1/2/rlk7*^AEQ^, similar to abolished CEP4-triggered calcium influx and MAPK activation in this mutant background (Supplementary Fig. 10B, Fig. 3E, F). Unlike *cepr1-3/2-4* (Supplementary Fig. 7A), the triple receptor mutant did not show elevated basal ethylene levels, raising the question whether RLK7 promotes ethylene accumulation in the absence of CEPR1/CEPR2 (Supplementary Fig. 10B). Also, flg22-induced MAPK activation and/or *FRK1* expression were unaltered in *cep6x* and *cepr1/2/rlk7*^AEQ^ (Supplementary Fig. 10C-E).

**Figure 5:**
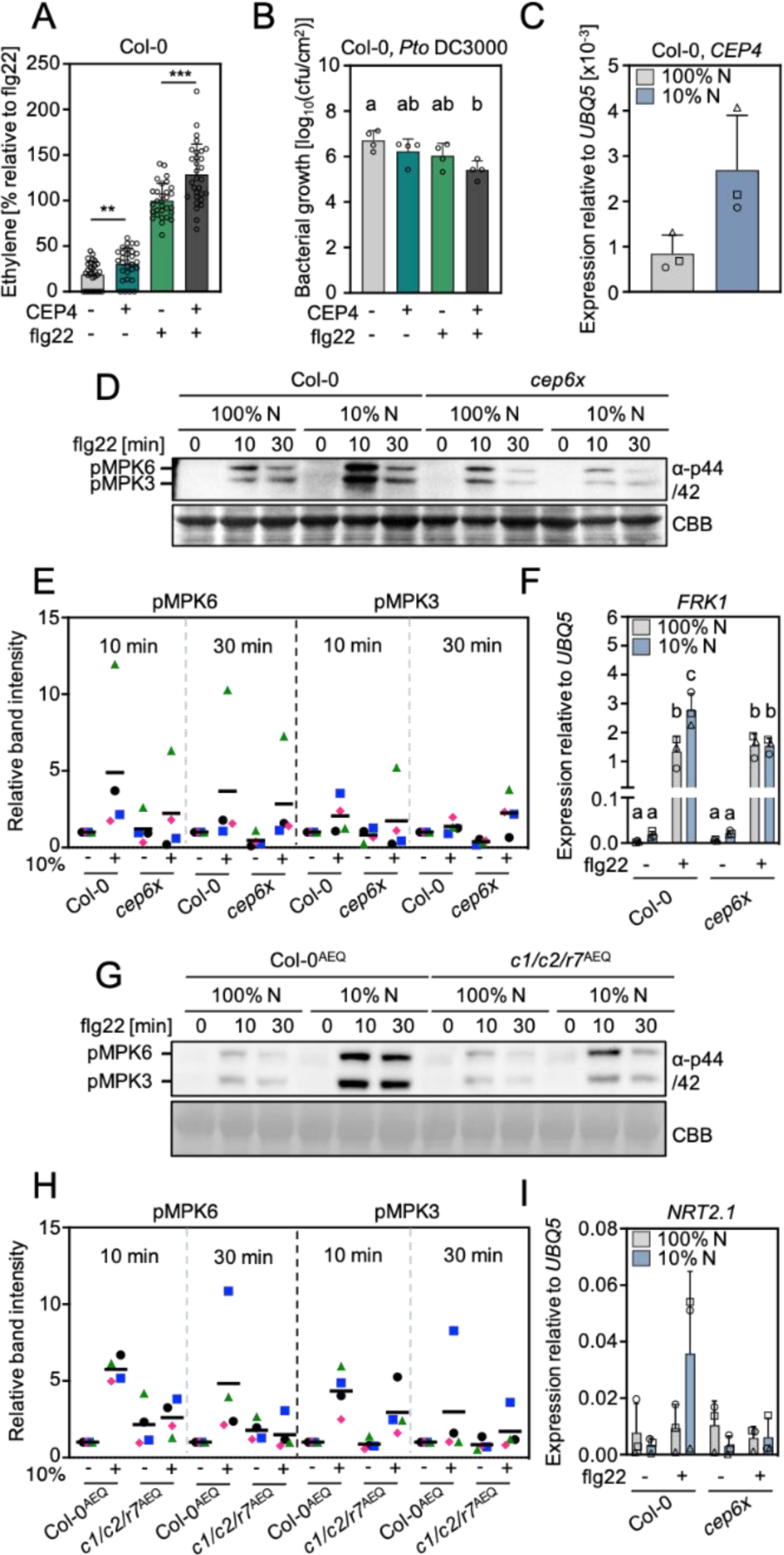
CEPs promote FLS2 signaling during reduced N conditions. A) Ethylene concentration in Col-0 leaf discs 3.5 h after mock (ddH_2_O), CEP4 (1 μM) and/or flg22 (500 nM) treatment; n = 30, seven pooled experiments ± SD (two-tailed Student’s t-test, ** p<0.05, *** p = 0.0001). **B)** cfu of *Pto* DC3000 (3 dpi) in Col-0 leaves upon mock (ddH_2_O), CEP4 (1 μM) and/or flg22 (100 nM) pre-treatment; n = 4 ± SD (one-way ANOVA, Tukey post hoc, a-b p<0.05). **C)** RT-qPCR of *CEP4* after 24 h transfer of Col-0 seedlings to 100% N or 10 % N medium. Housekeeping gene *UBQ5;* n = 3 ± SD. **D)** MAPK activation upon flg22 (100 nM) treatment after 24 h transfer of seedlings to 100% N or 10% N medium. Western blots were probed with α-p44/42. CBB = Coomassie brilliant blue. **E)** Quantification of pMPK6/MPK3 band intensities normalized to the CBB band and relative to 100% N flg22 set as 1 using ImageJ software; n = 4 biological replicates. **F)** RT-qPCR of *FRK1* upon flg22 (100 nM) treatment for 4 h after 24 h transfer of seedlings to 100% N or 10% N medium. Housekeeping gene *UBQ5*; n = 3 ± SD (one-way ANOVA, Tukey post hoc, a-b p<0.01, b-c ≤ 0,001, a-c <0.0001). **F)** MAPK activation upon flg22 (100 nM) treatment after 24 h transfer of seedlings to normal N or reduced N medium. Western blots were probed with α-p44/42. CBB = Coomassie brilliant blue. **H)** Quantification of pMPK6/MPK3 band intensities normalized to the CBB band and relative to 100% N flg22 set as 1 using ImageJ software.; n = 4 biological replicates. **I)** RT-qPCR of *NRT2.1* upon flg22 (100 nM) treatment of seedlings for 4 h after 24 h transfer to 100% N or 10% N. Housekeeping gene *UBQ5*; n = 3 ± SD. Different symbols in figures **C**, **E**, **F**, **H** and **I** represent independent experiments. All experiments were repeated at least three times in independent biological repeats with similar results.

As modulators of systemic N-demand signaling, several *CEPs* are transcriptionally upregulated in N-starved roots, including *CEP1* (*22*, *55*, *56*). It is known that the N status of plants affects disease resistance to different pathogens, but the underlying molecular mechanisms remain unknown (*57*, *58*). We noticed that *CEP4* is transcriptionally upregulated upon placing seedlings in ½ MS containing 10% N concentrations compared to the control medium (100% N ½ MS medium, 20 mM NO_3_^-^, 10 mM NH_4_^+^, Fig. 5C). Importantly, N starvation promoted resistance to *Pto* (*59*), suggesting a connection between N homeostasis and antibacterial resistance. We hypothesized that CEPs coordinate the plant’s N status with cell surface immunity. We transferred two-week-old seedlings for 24 h to ½ MS medium with different N concentrations before challenging them with flg22. Ten percent N and 5% N medium promoted flg22-induced MAPK activation, while 1% N did not (Supplementary Fig. 10F). This suggests that different N concentrations modulate the ability to mount FLS2 signaling. The promotion of flg22-induced MAPK activation upon 10% N treatment was compromised in *cep6x*, suggesting that CEPs promote FLS2 signaling under reduced N conditions (Fig. 5D, E). We confirmed this phenotype by measuring *FRK1* expression upon 10% N treatment and subsequent flg22 elicitation in wild-type and *cep6x* (Fig. 5F). We next tested whether CEP receptors are required for the N-dependent regulation of FLS2 signaling. Indeed, 10% N-promoted flg22-induced MAPK activation was reduced in *cepr1/2/rlk7*^AEQ^ (Fig. 5G, H).

We raised the question whether the impairment in FLS2 signaling under 10% N conditions in *cep6x* and *cepr1/2/rlk7*^AEQ^ mutants might be associated with defects in N homeostasis. Indeed, the No-0 *cepr1/2* mutant has paler leaves, smaller rosette size and constitutive anthocyanin accumulation, related to defects in nitrate uptake (*22*) but the *cep6x* shoot is indistinguishable from the wild-type plant (Supplementary Fig. 2D). We grew *cep6x* and *cepr1/2/rlk7*^AEQ^ seedlings in 100% N, 10% N and 1% N medium. We noticed a mild reduction in seedling growth of all genotypes in 10% N medium, which was more pronounced in 1% N medium after seven days (Supplementary Fig. 10G, H). The *cep6x* mutant phenotype was indistinguishable from the WT (Supplementary Fig. 10G). The *cepr1/2/rlk7*^AEQ^ seedlings also showed similar growth at lower N concentrations compared to its WT control (Supplementary Fig. 10H). This suggests that 10% N-induced promotion of flg22-triggered responses is not a mere consequence of deregulated metabolism in *cep6x* and *cepr1/2/rlk7*^AEQ^. Strongly reduced nitrate concentrations enhance the expression of the high-affinity nitrate transporter *NRT2.1* (*60–62*). We could not detect enhanced *NRT2.1* expression in WT under 10% N conditions, which remains higher than nitrate concentrations previously tested for *NRT2.1* expression (2 mM nitrate vs 1 mM) (Fig. 5I) (*61*). However, flg22 promoted *NRT2.1* transcript accumulation which was abrogated in *cep6x* (Fig. 5I). Altogether, these results indicate a CEP-and CEP receptor-dependent connection between FLS2-triggered PTI and the plant’s N supply, revealing a previously unknown mechanism of signaling cross-talk between cell surface immunity and the plant’s nutritional status.

## Discussion

This work shows that CEP perception in Arabidopsis is achieved by three partially cell type and tissue-specific receptors that regulate plant immunity and likely coordinate biotic stress with plant nutritional cues. CEP4 binds to both CEPR2 and RLK7, and CEP4 outputs show variable grades of dependency on specific receptors. The canonical group I CEP1 and CEP9.5, however, exclusively depend on CEPR1/2 and CEP1 does not bind to RLK7. The overlap of *CEP4* with *RLK7* expression in leaves, but not with *CEPR1* and *CEPR2*, suggests a combination of cell autonomous and short-distance signaling contributing to CEP-mediated immune modulation. Our work reveals an unexpected and previously undescribed complexity of phytocytokine signaling. A challenge for the future will be to determine how three CEP receptors with distinct expression patterns integrate responses between tissues and the concerted action of multiple ligands. RLK7 also recognizes PIPs to regulate growth, salt stress and immunity (*50*, *52*) (Fig. 3J). RLK7 binds PIP1 with higher affinity and likely preferentially recognizes this ligand. Members of the CLE peptide family also bind multiple receptors with variable affinities to regulate epidermal cell patterning (*44*, *63*). Dissecting spatiotemporal ligand availability and CEP4-PIP1 signaling specificity will be an important future task. RLK7 binds CEP4, but not CEP1, suggesting that CEP4’s receptor specificities are unique among CEPs, likely caused by its distinct sequence (Supplementary Fig. 1D). It will be interesting to compare the molecular mechanisms of PIP1/CEP4-RLK7 and CEP1/CEP4-CEPR2/CEPR1 recognition and activation in future structure analyses. Plant peptides may be versatile tools for the evolution of plant plasticity. High numbers of diverse ligands and receptors may provide the plant with vast combinations for fine-tuned signal outputs during adaption to environmental challenges.

Our data suggest that short-term reduction in N supplementation promotes *CEP* expression to promote FLS2 activation. Other phytocytokines modulate immune signaling by regulating PRR abundance or receptor complex formation and dynamics, but in most cases the mechanism remains unknown (*7–9*, *11*). It will be interesting to reveal whether CEPs directly or indirectly modulate PRR signaling.

Accumulating evidence suggests a direct integration of nutrient homeostasis and PTI in plants. Perception of flg22 by FLS2 induces PHT1.4 phosphorylation to inhibit phosphate (Pi) uptake and promote root immunity (*64*). It remains unknown whether immune activation also regulates N transport in root or shoot tissue. CEPs induce nitrate, Pi and sulfate uptake, suggesting CEP-CEPR1/CEPR2/RLK7-dependent modulation of several transporter pathways (*65*). This raises the question whether nutrient uptake directly contributes to CEP-mediated immune modulation. The main source of inorganic N for plant utilization is nitrate, which also functions as a signaling molecule to induce adaptive growth responses (*66*). Nitrate is sensed by the plasma membrane transceptor NRT1.1 and the nuclear transcriptional regulator NLP7 (*67*, *68*). NRT1.1 and similar transporters are regulated by phosphorylation to control transport activity, including NRT1.2 phosphorylation by CEPR2 (*67*, *69*– *73*), which we identified as a CEP4 receptor. It will be interesting to resolve whether and how cell surface signaling and nitrogen sensing/transport directly or indirectly intersect. Since seedlings grown under high N concentrations (as provided by ½ MS medium) show reduced flg22 responsiveness, it is also possible that N saturation inhibits PTI by suppressing *CEP* expression and accumulation.

CEPR1/CEPR2/RLK7 and CEPs are widely conserved among angiosperms, including crop plants (*40*). CEPs are important to promote nodulation in legumes and also regulate sucrose-dependent root growth inhibition and fecundity (*28*, *41*, *74*, *75*). This places CEPs as central integrators of biotic interactions (symbiosis and pathogen defense) with plant nutrition, growth and development (*65*, *76*). It will be critical to understand whether N-dependent and CEP-mediated PTI modulation extends beyond Arabidopsis, and to decipher how CEPs can promote immunity and control symbiosis in diverse species. This will provide important insights for future crop improvement strategies that coordinate crop nutrition with disease resistance.

## Materials and methods

### Molecular cloning

To generate *CEP4* overexpression lines, the coding sequence of *CEP4* (AT2G35612) was synthesized (Twist Bioscience, USA) with attB attachment sites for subsequent gateway cloning into pDONRZeo (Invitrogen, USA) and recombination with pB7WG2 (VIB, Ghent). To generate CRISPR-Cas9 mutants, appropriate target sites (two per gene of interest) were designed using the software tool chopchop (https://chopchop.cbu.uib.no/). Individual guide RNA constructs containing gene-specific target sites were synthesized (Twist Bioscience, USA) and subsequently stacked in a GoldenGate-adapted pUC18-based vector. To generate different order CRISPR *cep* mutants, *cepr1/2*^AEQ^, *rlk7/iku2*^AEQ^ and *rlk7*^AEQ^ 12, 4, 4 and 2 target site-containing gRNA constructs were stacked, respectively (Supplementary Table 2). Together with FastRed-pRPS5::Cas9, higher-order gRNA stacks were subsequently cloned into pICSL4723 for in planta expression (*77*).

To generate the *pCEP4::NLS-3xmVenus*, *pCEPR1::NLS-3xmVenus* and *pCEPR2::NLS-3xmVenus* reporter constructs, 1000, 1696 and 2788 bp fragments upstream of the start codon, respectively, were amplified from genomic DNA and assembled together with the sequence coding for the nuclear localization signal of SV40 large T antigen followed by 3 consecutive mVenus YFP fluorophores (*78*) into a GoldenGate-modified pCB302 binary vector for plant expression. For *pRLK7* the 1957 bp promoter sequence upstream from the start codon was amplified with primers containing attB attachment sites for subsequent gateway cloning into pDONRZeo (Invitrogen, USA) and recombination with promotor::Venus (YFP)-H2B destination vector (*78*). For *CEPR1* (AT5G49660) and *CEPR2* (AT1G72180) overexpression lines, the coding sequence of both genes was amplified from cDNA with attB attachment sites for subsequent cloning into a pDONR223 (Invitrogen, USA) and recombination with pK7FWG2 (VIB Ghent, Belgium). All of the generated plant expression constructs were subsequently transformed into *Agrobacterium tumefaciens* strain GV3101 before floral dip transformation of Arabidopsis. All primers used for cloning are listed in (Supplementary Table 3).

### Plant material and growth conditions

Arabidopsis Col-0, Col-0^AEQ^ (*36*) and No-0 were used as wild types for experiments and generation of transgenic lines or CRISPR mutants. The *cepr1-3* (GK-467C01), *cepr2-3* (SALK_014533), *rlk7-1* (SALK_056583), *rlk7-3* (SALK_120595), *iku2-4* (Salk_073260) and the novel *cepr2-4* allele (GK-695D11) were obtained from NASC (UK) (*28*, *52*, *79*, *80*). The No-0 *cepr2-1xcepr2-1* was obtained from RIKEN (Japan) (*22*). T-DNA insertion mutants were genotyped by PCR using T-DNA- and gene-specific primers as listed in Supplementary Table 3. The lack of *CEPR2* transcript in *cepr2-4* (GK-695D11) was determined by semi-quantitative PCR from cDNA using *cepr2-4* genotyping primers (Supplementary Fig. 5A). The *cepr1-3/2-4* double mutant was obtained by genetic crossing. The *bak1-5/bkk1* mutant was characterized previously (*81*). For visualizing tissue-specific *FRK1* expression, a *pFRK1::NLS-3xmVenus* line was used (*82*). To isolate homozygous CRISPR mutants, pICSL4723 transformant T1 seeds showing red fluorescence were selected and grown on soil before genotyping with gene-specific primers and Sanger sequencing (Supplementary Table 3). Mutants lacking the transgene were identified by loss of fluorescence. To generate the *cep6x 35S::CEP4* lines, the same pB7WG2 CEP4 construct used for the generation of *CEP4* overexpression lines was utilized for floral dip transformation of homozygous *cep6x*.

Plants for physiological assays involving mature plants were vernalized for 2-3 days in the dark at 4°C and later grown in individual pots in environmentally controlled growth rooms (20-21°C, 55% relative humidity, 8 h photoperiod). For seedling-based assays, seeds were sterilized using chlorine gas and grown axenically on ½ Murashige and Skoog (MS) media supplemented with vitamins (Duchefa, Netherlands), 1% sucrose, with or without 0.8% agarose at 22°C and a 16 h photoperiod unless stated otherwise. For experiments using ½ MS medium with reduced N concentrations, modified MS salts without nitrogen-containing compounds (Duchefa, Netherlands) were used and supplemented with KNO_3_/NH_4_NO_3_ to achieve 100% N (KNO_3_ 9.395 mM, NH_4_NO_3_ 10.305 mM), 10% N (KNO_3_ 0.9395 mM, NH_4_NO_3_ 1.0305 mM), 5% (KNO_3_ 0.4698 mM, NH_4_NO_3_ 0.5153 mM) and 1% (KNO_3_ 0.09395 mM, NH_4_NO_3_ 0.10305 mM) conditions. To keep the ionic strength equal in 10%, 5% and 1% N conditions, media were supplemented with 90% (8.455 mM), 95% (8.925 mM) and 99% (9.301 mM) KCl, respectively.

### Grafting

Arabidopsis seedlings were grown vertically on ½ MS agar medium without sucrose in short-day conditions seven days before grafting. Grafting was performed aseptically under a stereo microscope as previously described (*83*). Vertically mounted plates with reciprocally grafted seedlings were returned to short-day conditions for 10 days. Healthy seedlings were transferred to the soil.

### Imaging and microscopy

Confocal laser-scanning microscopy was performed using a Leica TCS SP5 (Leica, Germany) microscope (with Leica Application Suite X 3.7.4.23463) and all the pictures were taken with a 20x water immersion objective. For the mVenus fluorophore, pictures were imaged with argon laser excitation at 514nm and a detection window of 525-535 nm. Propidium iodide was visualized using DPSS 561 laser emitting at 561 nm with a detection window of 575-590 nm. For imagining the expression pattern of NLS-3xmVenus or Venus (YFP)-H2B under the control of different promoters (*pCEP4*, *pCEPR1*, *pCEPR2* and *pRLK7*), vertically-grown seedlings were stained with propidium iodide immediately before microscopic analysis. For imagining the *pFRK1::NLS-3xmVenus* reporter line, 12-day-old seedlings were transferred to a 24-well-plate containing ddH_2_O with or without (mock) peptides in the indicated concentrations. For comparison of *FRK1* promoter activity, seedlings were analysed by confocal microscopy using identical laser intensities and interval/number of slices for Z stack projection 16 h after treatment.

### Calcium influx assay

Apoaequorin-expressing liquid-grown eight-day-old seedlings were transferred individually to a 96-well plate containing 100 µl of 5 µM coelenterazine-h (PJK Biotech, Germany) and incubated in the dark overnight. Luminescence was measured using a plate reader (Luminoskan Ascent 2.1, Thermo Fisher Scientific, USA). Background luminescence was recorded by scanning each well 12 times at 10 s intervals, before adding a 25 µl elicitor solution to the indicated final concentration. Luminescence was recorded for 30 min at the same interval. The remaining aequorin was discharged using 2 M CaCl_2_, 20% ethanol. The values for cytosolic Ca^2+^ concentrations ([Ca^2+^]_cyt_) were calculated as luminescence counts per second relative to total luminescence counts remaining (L/L_max_).

### Ethylene measurement

Leaf discs (4 mm in diameter) from four-to-five-week-old soil-grown Arabidopsis were recovered overnight in ddH_2_O. Three leaf discs per sample were transferred to a glass vial containing 500 µl of ddH_2_O before adding ddH_2_O (mock) or peptides to the indicated final concentration. Glass vials were capped with a rubber lid and incubated under gentle agitation for 3.5 h. One mL of the vial headspace was extracted with a syringe and injected into a Varian 3300 gas chromatograph (Varian, USA) to measure ethylene.

### MAPK activation and western blot analysis

Five-day old Arabidopsis seedlings growing on ½ MS agar plates, were transferred into a 24-well plate containing liquid medium for seven days. 24 h before the experiment, seedlings were equilibrated in a fresh ½ MS medium. For N reduction experiments, modified ½ MS containing 100% N, 10% N, 5% N and 1 % N supplemented with KCl was used. MAPK activation was elicited by adding the peptides to the indicated concentrations. Six seedlings per sample were harvested, frozen in liquid nitrogen and homogenized using a tissue lyser (Qiagen, Germany). Proteins were extracted using a buffer containing 50 mM Tris-HCl (pH 7.5), 50 mM NaCl, 10% glycerol, 5 mM DTT, 1% protease inhibitor cocktail, 1 mM phenylmethylsulfonyl fluoride (PMSF), 1% IGEPAL, 10 mM EGTA, 2 mM NaF, 2 mM Na_3_VO_4_, 2 mM Na_2_MoO_4_, 15 mM ß-Glycerophosphate and 15 mM p-nitrophenylphosphate before analysis by SDS-PAGE and western blot. Phosphorylated MAPKs were detected by α-p44/42 antibodies (Cell Signaling, USA). To quantify the intensity of the specific bands, ImageJ software (version 1.53t) was used. Each band was selected with the same-sized frame and the intensity peak was determined. The area under each peak was calculated and normalized to Coomassie staining as a measure of relative band intensity (RBI). The RBI of the wild-type genotypes at 100% N upon flg22 treatment was set to one.

To determine CEPR1 and CEPR2 protein levels in *35S::CEPR1-GFP* and *35S::CEPR2-GFP* overexpression lines, seedlings were grown in ½ MS liquid medium for 12 days. Afterwards, harvested seedlings were frozen in liquid nitrogen, homogenized using a tissue lyser (Qiagen, Germany) and the proteins were isolated using an extraction buffer contenting 50 mM Tris-HCl (pH 7.5), 50 mM NaCl, 10% glycerol, 2 mM EDTA, 2 mM DTT, 1% protease inhibitor cocktail, 1 mM phenylmethylsulfonyl fluoride (PMSF) and 1% IGEPAL. After SDS-PAGE and western blot, GFP-tagged proteins were detected by α-GFP antibodies (ChromoTek).

### Seedling growth inhibition

Arabidopsis seedlings were grown for five days on ½ MS agar plates before the transfer of individual seedlings into each well of a 48-well plate containing liquid medium with or without elicitors in the indicated concentration. After seven-day treatment, the fresh weight of individual seedlings was measured.

### Pathogen growth assay

*Pseudomonas syringae* pv. *tomato* (*Pto)* DC3000 and *Pto* lacking the effector molecular coronatine *Pto*^COR-^ were grown on King’s B agar plates containing 50 μg/mL rifampicin and 50 μg/mL kanamycin at 28°C. After two-to-three days bacteria were resuspended in ddH_2_O containing 0.04% Silwet L77 (Sigma Aldrich, USA). The bacterial suspension was adjusted to an OD_600_ = 0.2 (10^8^ cfu/mL) for *Pto*^COR*-*^ or OD_600_ = 0.02 (10^7^ cfu/mL) for *Pto* DC3000 before spray inoculating four-to-five-week-old plants. For peptide-induced resistance in local tissues, ddH_2_O (mock), flg22 (100 nM) and/or CEP4 (1 µM) were syringe-infiltrated into mature leaves. After 24 h, *Pto* DC3000 (OD_600_ = 0.0002, 10^5^ cfu/mL) was syringe-infiltrated into pre-treated leaves and incubated for three days before determining bacterial counts. For CEP4-induced resistance in systemic tissues, CEP4 (1 or 5 µM) or ddH_2_O (mock) were syringe-infiltrated into the first two true leaves of young three-to-four-week-old Arabidopsis. After four days, *Pto* DC3000 (OD_600_=0.0002, 10^5^ cfu/mL) was syringe-infiltrated into leaves three and four of the pre-treated plants. Bacterial counts were determined four days after infection.

### Systemic acquired resistance

SAR experiments were performed as previously described (*84*). Briefly, plants were cultivated in a mixture of substrate (Floradur) and silica sand in a 5:1 ratio under short day (SD) conditions (10 h) in a growth chamber at 22 °C /18 °C (day/night) with a light intensity of 100 μmol m^-2^s^-1^, and 70% relative humidity (RH). SAR assays were performed using *Pto* DC3000 and *Pto* AvrRpm1. Bacteria were grown on NYGA media (0.5% peptone, 0.3% yeast extract, 2% glycerol, 1.8% agar, 50 µg/mL kanamycin, 50 µg/mL Rifampicin) at 28° C before infiltration. Freshly grown *Pto* avrRpm1 was diluted in 10 mM MgCl_2_ (to reach a final concentration of 1×10^6^ cfu/mL) and syringe-infiltrated in the first two true leaves of four-and-a-half-week-old plants. Concurrently, 10 mM MgCl_2_ was applied to a separate set of plants as the mock control treatment. Three days after *Pto* avrRpm1 infiltration, plants were challenged in their 3^rd^ and 4th leaves with *Pto* DC3000 (1×10^5^ cfu/mL). Bacterial titers were determined four days after *Pto* DC3000 infection.

### Gene expression analysis

For seedlings-based assays, 12-day-old liquid-grown seedlings were equilibrated in fresh medium for 24 h before treatment with the indicated peptides. For adult plants, four-to-five-week-old Arabidopsis leaves were syringe-infiltrated with ddH_2_O (mock), flg22 (1 µM) or *Pto* DC3000 (OD_600_ = 0.001, 5×10^5^cfu/mL) and incubated for 24 h. All samples for RT-qPCR analysis were harvested at the indicated time points, frozen in liquid nitrogen and homogenized using a tissue lyser (Qiagen, Germany). Total RNA was isolated using TRIzol reagent (Roche, Switzerland) and purified using Direct-zol™ RNA Miniprep Plus kit (Zymo Research, Germany). 2 µg of the total RNA was digested with DNase I and reverse transcribed with oligo (dT)18 and Revert Aid reverse transcriptase. RT-qPCR experiments were performed using Takyon^TM^ Low ROX SYBR MasterMix (Eurogentec, Belgium) with the AriaMx Real-Time PCR system (Agilent Technologies, USA). Expression levels of all tested genes were normalized to the house-keeping gene *Ubiquitin 5* (*UBQ5*). Sequences of all primers used for RT-qPCR analysis are found in Supplementary Table 3.

### Expression and purification of recombinant receptor ectodomains

*Spodoptera frugiperda* codon-optimized synthetic genes (Invitrogen GeneArt), coding for Arabidopsis CEPR1 (residues 23 to 592), CEPR2 (residues 32 to 620) and RLK7 (residues 29 to 608) were cloned into a modified pFastBAC vector (Geneva Biotech) providing a 30K signal peptide (*85*), a C-terminal TEV (tobacco etch virus protease) cleavable site and a StrepII-9xHis affinity tag. For protein expression, *Trichoplusia ni Tnao38* cells (*86*) were infected with CEPR1, CEPR2 or RLK7 virus with a multiplicity of infection (MOI) of 3 and incubated one day at 28°C and two days at 21°C at 110 rounds per minute (rpm). The secreted proteins were purified by Ni^2+^ (HisTrap excel, Cytiva, equilibrated in 25 mM KP_i_ pH 7.8 and 500 mM NaCl) followed by Strep (Strep-Tactin Superflow high-capacity, IBA Lifesciences, equilibrated in 25 mM Tris pH 8.0, 250 mM NaCl, 1 mM EDTA) affinity chromatography. All proteins were incubated with TEV protease to remove the tags. Proteins were purified by SEC on a Superdex 200 Increase 10/300 GL column (Cytiva, USA) equilibrated in 20 mM citrate pH 5.0, 150 mM NaCl and further concentrated using Amicon Ultra concentrators from Millipore (Merck, Germany) with a 30,000 Da molecular weight cut-off. Purity and structural integrity of the different proteins were assessed by SDS-PAGE.

### Analytical size-exclusion (SEC) chromatography

Analytical SEC experiments were performed using a Superdex 200 Increase 10/300 GL column (GE, USA). The columns were pre-equilibrated in 20 mM citric acid pH 5, 150 mM NaCl. 150 μg of CEPR1, CEPR2 and RLK7 were injected sequentially onto the column and eluted at 0.5 mL/min. Ultraviolet absorbance (UV) at 280 nm was used to monitor the elution of the proteins. The peak fractions were analyzed by SDS-PAGE followed by Coomassie blue staining.

### Isothermal titration calorimetry (ITC)

Experiments were performed at 25°C on a MicroCal PEAQ-ITC (Malvern Instruments, UK) using a 200 µL standard cell and a 40 μL titration syringe. CEP1, CEP4, CEP4^scr^ and PIP1 peptides were dissolved in the SEC buffer to match the receptor protein. A typical experiment consisted of injecting 1 μL of a 300 μM solution of the peptide into 30 μM CEPR2 or RLK7 solution in the cell at 150 s intervals. ITC data were corrected for the heat of dilution by subtracting the mixing enthalpies for titrant solution injections into protein-free ITC buffer. Experiments were done in duplicates and data were analyzed using the MicroCal PEAQ-ITC Analysis Software provided by the manufacturer. The N values were fitted to 1 in the analysis.

### Synthetic peptides

The flg22 peptide was kindly provided by Dr. Justin Lee (IPB Halle). Other peptides were synthesized by Pepmic (China) with at least 90% purity and dissolved in ddH_2_O. Sequences of all the synthetic peptides can be found in Supplementary Table 1.

### Quantification and statistical analysis

Statistical analyses were performed using GraphPad Prism (Version 10.1.0). Sample size, p-values and statistical methods employed are described in the respective figure legends.

## Data and material availability

All data are available in the main text or the supplementary materials. All newly generated mutant lines are available upon request to M.S.

## Author contributions

Conceptualization: MS; Investigation: JR, HL, HKL, CB, SN, CW, JM, ZC, VOL, MS; Funding acquisition: MS, JA, RH, ACV, MAD; Project administration: MS; Supervision: MS, JR, ACV, JS, MAD, RH; Writing – original draft: MS, JR; Writing – review & editing: HL, HKL, CB, SN, CW, JM, VOL, MAD, ACV, RH, JS.

## Acknowledgments

We thank Stefanie Ranf for providing GoldenGate vectors for molecular cloning and Ulrich Hammes for advice in grafting experiments. This work was funded by the following agencies: Deutsche Forschungsgemeinschaft (DFG) grants STE2448/3-1 (MS and JR), STE2448/4-1 (MS and HL) and SFB924 TP B06 (ACV), the Technical University of Munich (MS, JR, RH, ZC, CW and JM), the Research Council of Norway, Grant 230849 (VOL), Australian Research Council, Grant DP200101885 (MAD), the University of Lausanne (CB), the Swiss National Science Foundation grants no. 310030_204526 and the European Research Council (ERC) grant agreement no. 716358 (JS, HKL).

## Supplementary Figures

**Supplementary Fig. 1:**
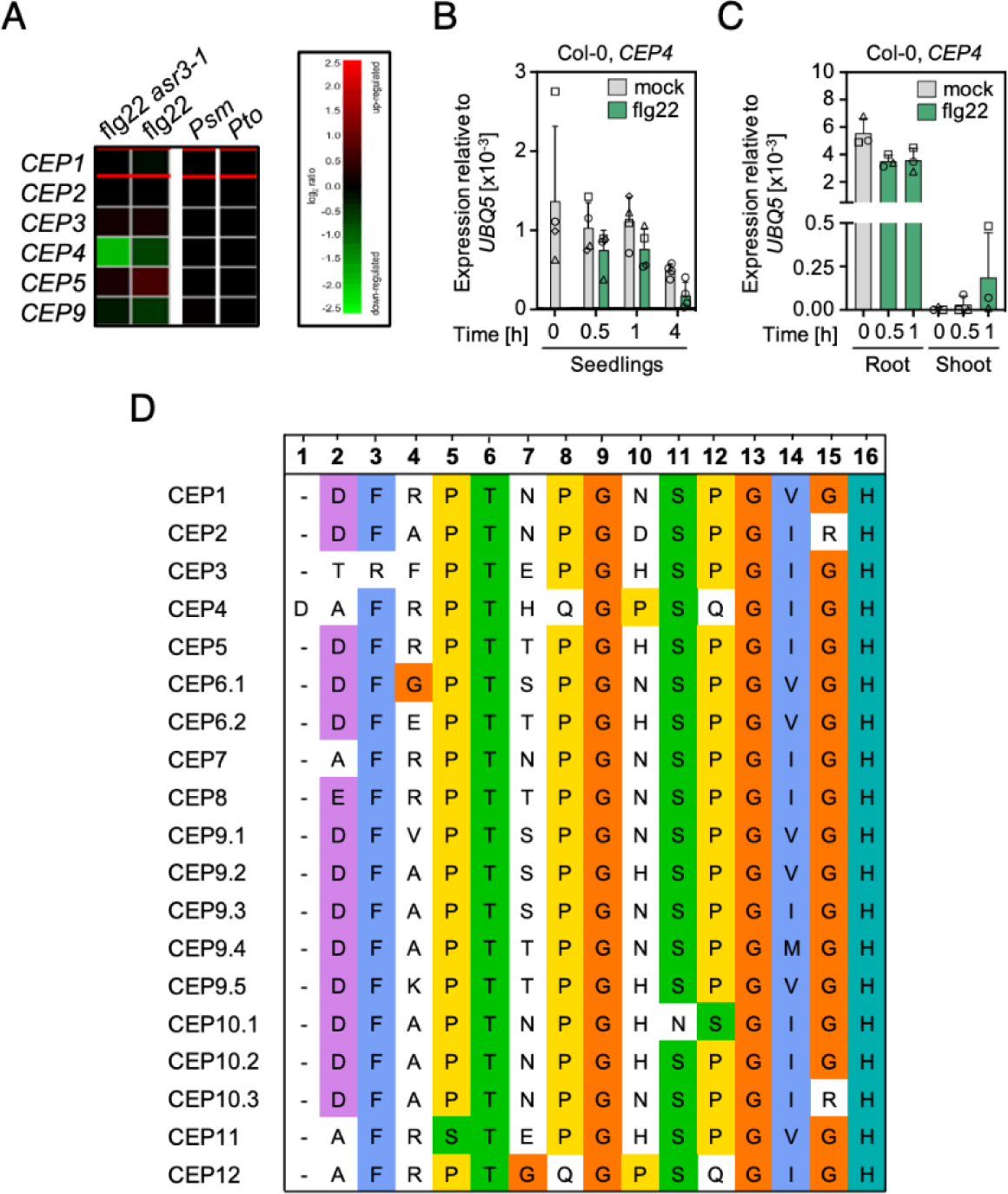
*CEP4* expression is differentially regulated after flg22 perception. **A)** *CEP4* is downregulated after flg22 treatment in an *asr3* mutant background (*32*). Data were obtained using Genevestigator software and are based on the AT_mRNASeq_ARABI_GL-1 data set. All *CEP* genes for which RNAseq data was available are depicted **B)** *CEP4* expression in seedlings is weakly downregulated after flg22 treatment. Col-0 seedlings were treated with flg22 (100 nM) or mock (ddH_2_O) for the indicated time, after which seedlings were harvested for RT-qPCR analysis of *CEP4* abundance. Housekeeping gene *UBQ5*; n = 3 ± SD. **C)** *CEP4* shows higher expression in the roots. Col-0 seedlings were treated with flg22 (100 nM) or mock (ddH_2_O) for the indicated time, after which seedlings were harvested for RT-qPCR analysis of *CEP4* abundance. Housekeeping gene *UBQ5*; n = 3 ± SD. **D)** Alignment of group I CEP peptide domains from *Arabidopsis thaliana* marked with colors based on amino acid properties and conservation. The conserved proline (P) residues are indicated in yellow. Different symbols in **B** and **C** represent independent biological experiments.

**Supplementary Fig. 2:**
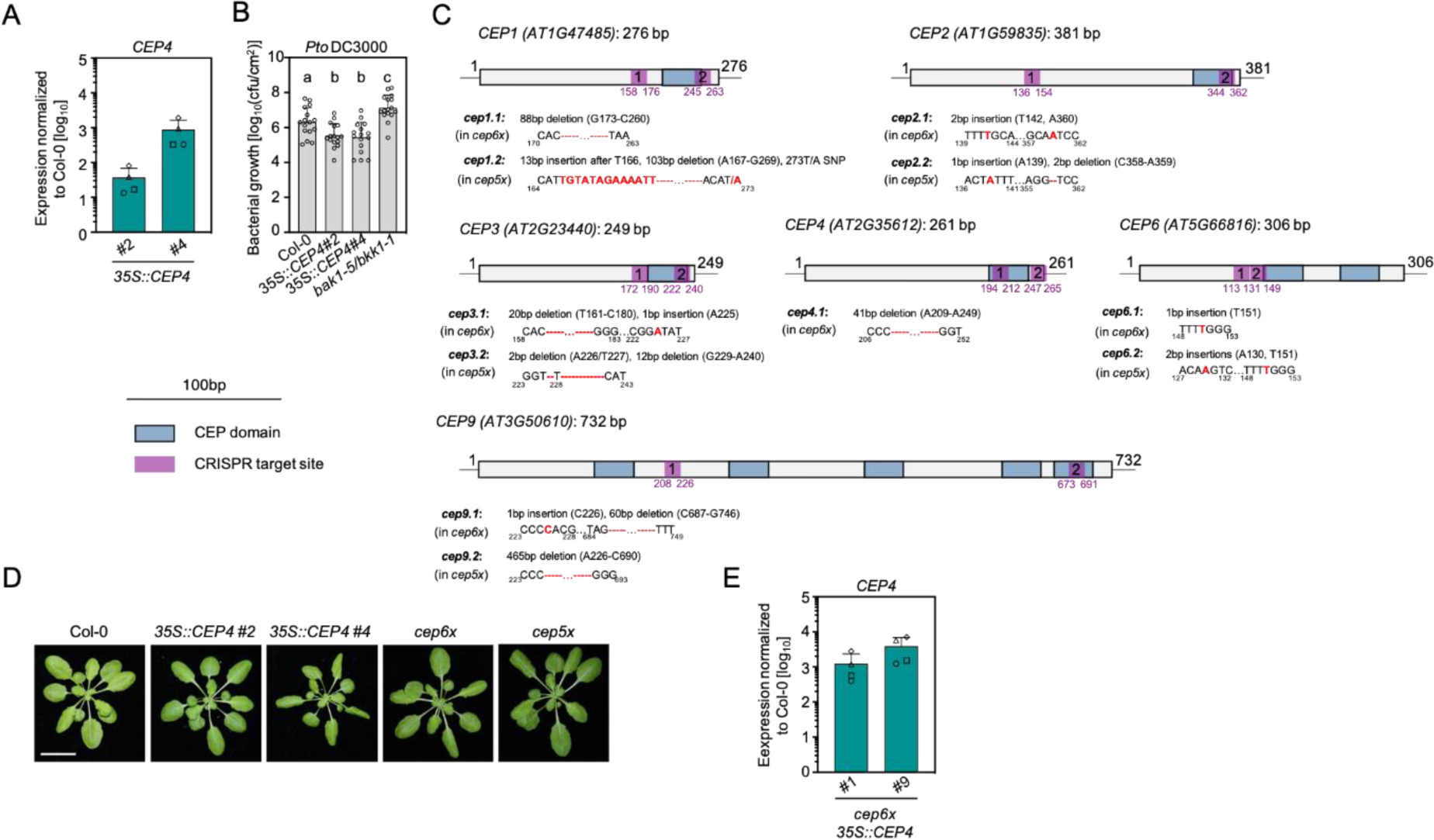
Characterization of *35S::CEP4*, CRISPR *cep* mutant alleles and *cep6x 35S::CEP4*. **A)** *CEP4* transcript levels in two independent *CEP4* overexpression lines, shown as fold induction compared to Col-0. Housekeeping gene *UBQ5*; n = 4 ± SD. **B)** Cfu of *Pto* DC3000 3 dpi upon spray infection; n = 16 pooled from four experiments ± SD (one-way ANOVA, Tukey post-hoc test; a-b/c p<0.05; b-c p<0.0001). **C)** Characterization of four independent *cep* mutant alleles. Schematic diagram of *CEP1*, *CEP2*, *CEP3*, *CEP4*, *CEP6* and *CEP9* gene structure and the CRISPR-Cas9-mediated mutation pattern detected by DNA sequencing. The locus number and the length of the coding sequence (CDS) are indicated above the scheme for each gene. CRISPR *cep6x* is mutated in *cep1/2/3/4/6/9* (alleles *cep1.1, cep2.1, cep3.1, cep4.1, cep6.1, cep9.1*), CRISPR *cep5x* is mutated in *cep1/2/3/6/9* with CEP4 wild-type (alleles *cep1.2, cep2.2, cep3.2, cep6.2, cep9.2*). The specific location and type of mutations for each gene are indicated in the schematics describing the mutants. The CEP domain is indicated in blue, and the two CRISPR target sites are indicated in purple. **D)** Pictures of 5-week-old plants of the indicated genotypes grown on soil; scale bar = 2 cm. **E)** *CEP4* transcript levels in two independent lines of *cep6x 35S::CEP4* shown as fold induction compared to Col-0; Housekeeping gene *UBQ5*; n = 4 ± SD. Different symbols in **A** and **E** represent independent biological repeats.

**Supplementary Fig. 3:**
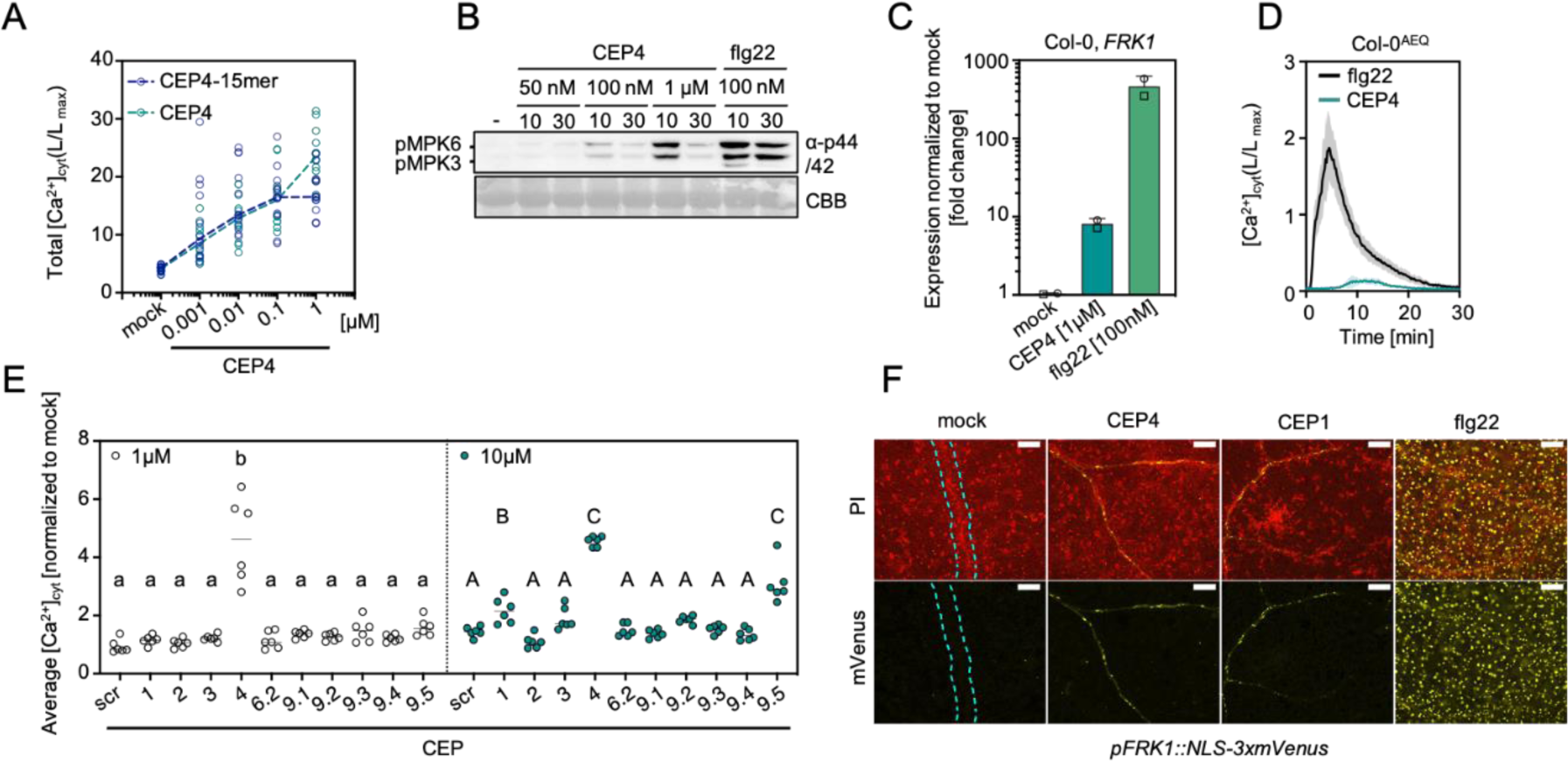
CEP4 is the strongest inducer of PTI responses on the whole tissue level. **A)** Both *CEP4* variants trigger influx of calcium ions. Col-0^AEQ^ seedlings were treated with the indicated concentrations of CEP4 and CEP4-15mer. ([Ca^2+^]_cyt_) was measured for 30 min. Shown is total calcium influx; n = 12 pooled from two experiments. **B)** MAPK activation in Col-0 upon treatment with indicated concentrations of CEP4 or flg22. Western blots were probed with α-p44/42. CBB = Coomassie brilliant blue. **C)** RT-qPCR of *FRK1* in seedlings upon mock (ddH_2_O), CEP4 (1 μM) or flg22 (100 nM) treatment for 4 h, shown as a fold induction compared to Col-0. Different symbols represent independent biological repeats. Housekeeping gene *UBQ5*; n = 2 ± SD. **D)** Kinetics of cytosolic calcium concentrations ([Ca^2+^]_cyt_) in Col-0^AEQ^ seedlings upon CEP4 (100 nM) and flg22 (100nM) treatment; n = 12, ± SD. **E)** Some CEPs induce Ca^2+^ influx at higher concentrations. Col-0^AEQ^ seedlings were treated with the indicated concentrations of CEPs. ([Ca^2+^]_cyt_) was measured for 30 min. The average calcium influx triggered by individual CEPs is normalized to mock (ddH_2_O). The dotted line represents different experiments. Statistical analysis was performed separately for each group. Statistical significance was compared to CEP4^scr^ of the indicated concentration; n = 6 (one-way ANOVA, Dunnett post hoc test, a-b p<0.0001, A-B p<0.001, A-C p<0.0001). **F)** CEP4 and CEP1 induce *FRK1* promoter activity in the vasculature. Representative images of NLS-3xmVenus signal in *pFRK1::NLS-3xmVenus* lines upon mock (ddH_2_O), CEP1 (100 nM), CEP4 (100 nM) or flg22 (100 nM) treatment for 16 h. Maximum projection of Z-stack for mVenus merged with Z-stacked propidium iodide (PI) signal. Cyan dotted line indicates vasculature,scale bar = 100 μm. The experiment was repeated three times in independent biological repeats with similar results.

**Supplementary Fig. 4:**
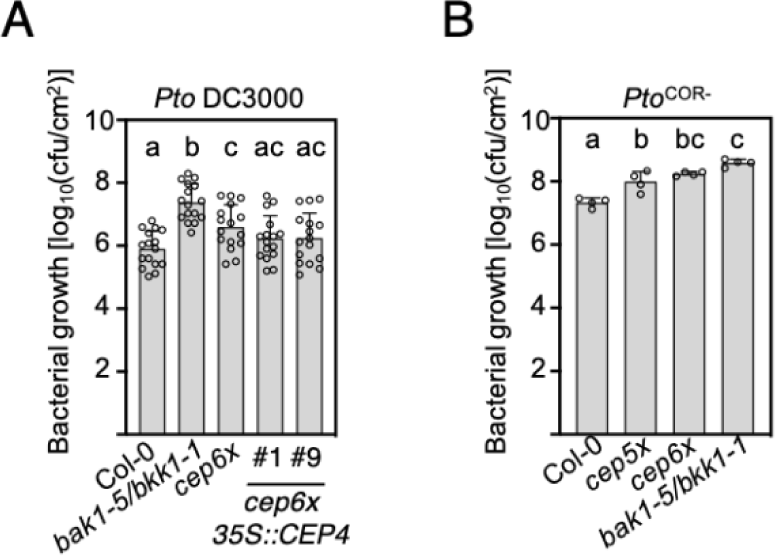
CEPs are important for resistance against *Pto*. **A)** *CEP4* overexpression does not fully rescue the CRISPR *cep6x* susceptibility to *Pto* DC3000. Cfu of *Pto* DC3000 (3 dpi) upon spray infection; n = 16 pooled from four experiments ± SD (one-way ANOVA, Tukey post-hoc test; a-b p<0.0001; a-c, b-c p<0.05; a/c-b p≤0.0001). **B)** Loss of *CEPs* increases Arabidopsis susceptibility to *Pto*^COR-^. Cfu of *Pto*^COR-^ (3 dpi) upon spray infection; n = 4 ± SD (one-way ANOVA, Tukey post-hoc test; a-b p = 0.0015; a-bc/a-c p<0.0001; b-c p = 0.0041). All experiments were repeated at least three times in independent biological repeats with similar results.

**Supplementary Fig. 5:**
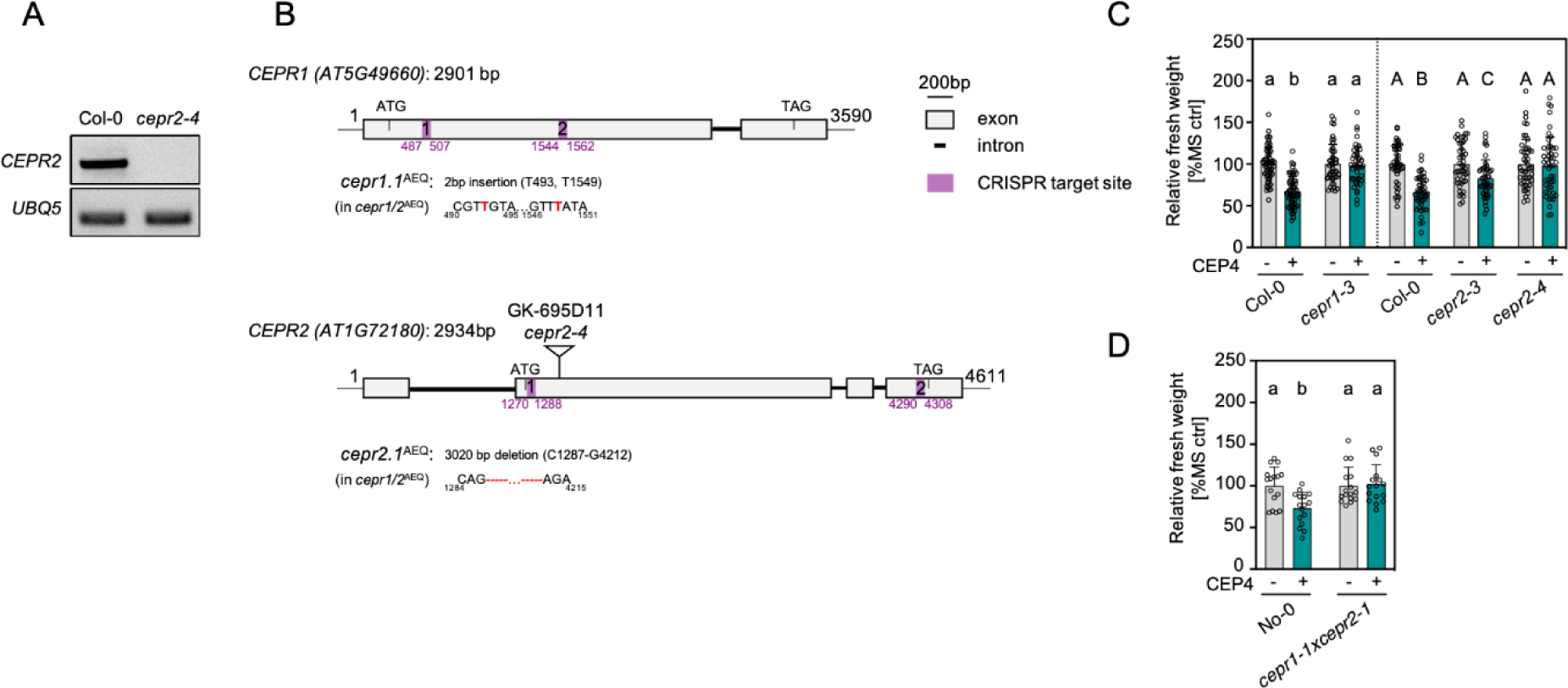
Characterization of *cepr2-4* and *cepr1/2*^AEQ^ mutants and CEP4-induced seedling growth inhibition. **A)** Characterization of the *cepr2-4* mutant. The *CEPR2* transcript could not be detected in *cepr2-4* mutant in a semi-quantitative RT-PCR analysis. Housekeeping gene *UBQ5*. **B)** Schematic diagram of the *CEPR1* and *CEPR2* genomic sequence, structure, and the CRISPR-Cas9-mediated mutation pattern detected by DNA sequencing. The locus number and the length of the coding sequence (CDS) are indicated above the scheme for each gene. The CRISPR *cepr1*/2^AEQ^ mutant was generated in Col-0^AEQ^ background. CRISPR *cepr1*^AEQ^ (in *cepr1/2*^AEQ^) has two 1 bp insertions, which lead to a frameshift mutation and an early stop codon. CRISPR *cepr2*^AEQ^ (in *cepr1/2*^AEQ^) has a 3020 bp deletion between the start and stop codon. The specific location and type of mutations for each gene are indicated in the schematics describing the mutants. The two CRISPR target sites are indicated in purple. The T-DNA insertion site in *cepr2-4* is indicated with a black triangle. **C)** CEPR1 and CEPR2 are similarly required for CEP4-induced seedling growth inhibition. Relative fresh weight of five-day-old seedlings treated with CEP4 (1 μM) for seven days. The dotted line represents different experiments. Statistical analysis was performed separately for each group; n = 42-60 and n = 46-48 pooled from five and four experiments, respectively (one–way ANOVA, Tukey post-hoc test, a-b p<0.0001; A-B p<0.0001; A/B-C p<0.05). **D)** CEP4-induced seedling growth inhibition is abolished in *cepr1-1xcepr2-1*. Relative fresh weight of five-day-old seedlings treated with CEP4 (1 μM) for seven days; n=16 ± SD (one-way ANOVA, Tukey post-hoc test, a-b p<0.01). Similar results were obtained in three independent biological repeats.

**Supplementary Fig. 6:**
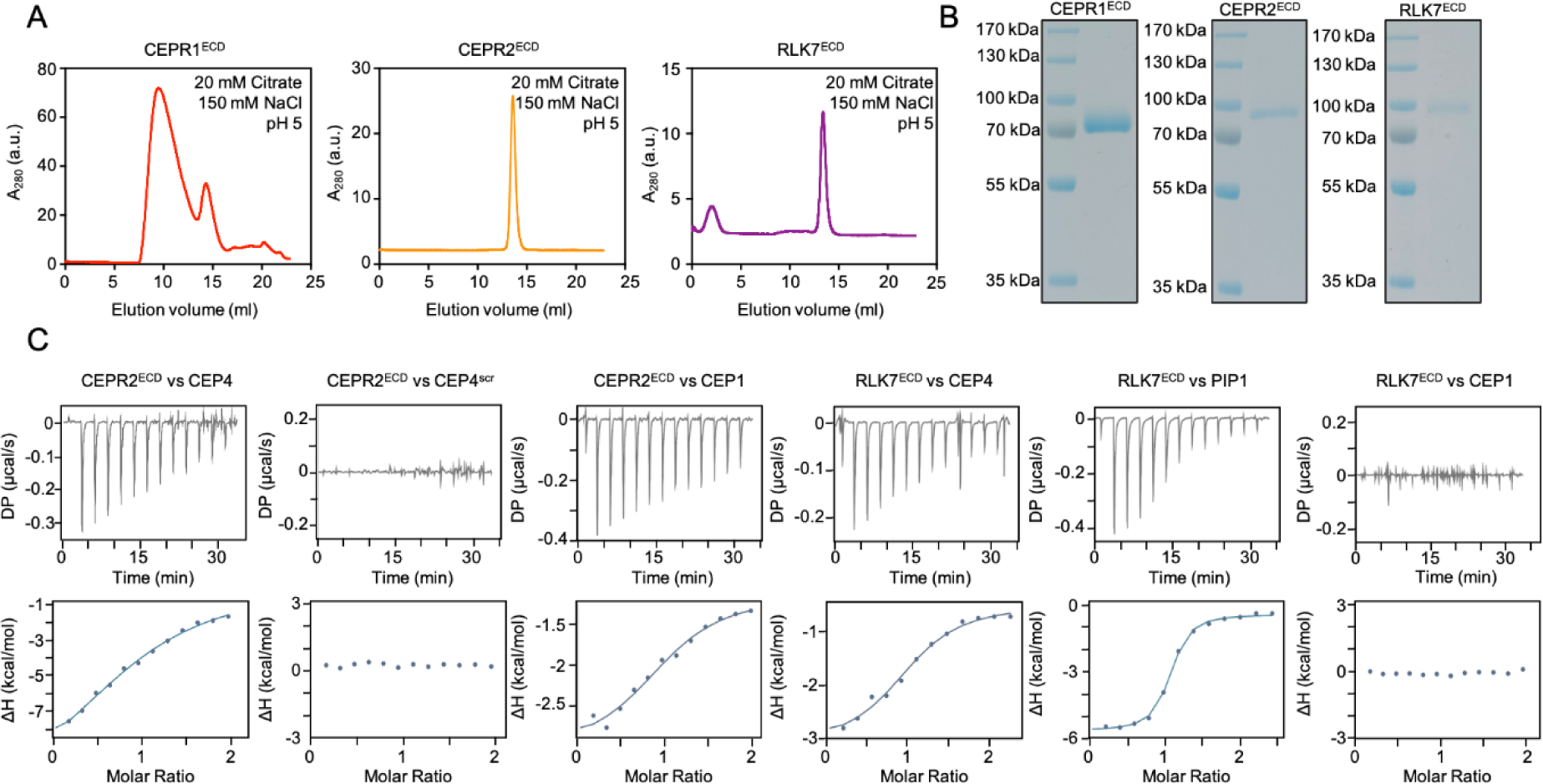
Quality control of recombinant CEPR1, CEPR2 and RLK7 ectodomains and supplemental ITC data. **A)** Analytical size-exclusion chromatography (SEC) experiments show CEPR1^ECD^ aggregates in comparison to single peaks for CEPR2^ECD^ and RLK7^ECD^. **B)** SDS-PAGE of the ECD peaks of the SEC analysis in **A**. **C)** ITC thermograms of the second technical repeat performed for each ectodomain and peptide analyzed in Fig. 2 (**D**-**F**) and Fig. 3 (**I**-**K**).

**Supplementary Fig. 7:**
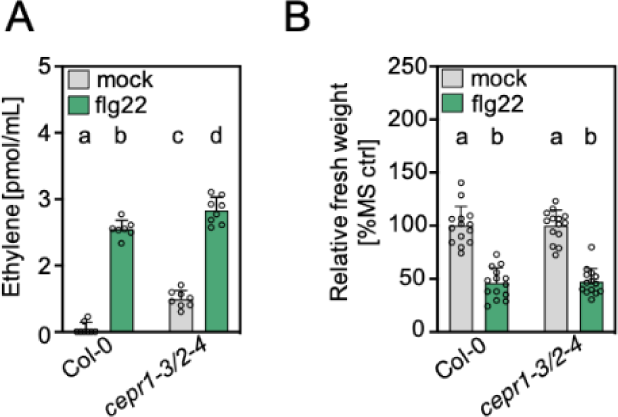
CEPR1 and CEPR2 are dispensable for flg22-induced ethylene accumulation and growth inhibition. **A)** Basal and flg22-triggered ethylene production is higher in *cepr1-3/2-4* mutants. Ethylene concentration in leaf discs upon mock (ddH_2_O) or CEP4 (1 μM) for 3.5 h; n = 7 -8 pooled from two experiments ± SD (one-way ANOVA, Tukey post-hoc test, a-b/c/d, c-b/d p<0.0001, b-d p<0.005). **B)** flg22-induced seedling growth inhibition is not affected in *cepr1-3/2-4*. Relative fresh weight of five-day-old seedlings treated with flg22 (100 nM) for seven days; n = 14 ± SD (one-way ANOVA, Tukey post-hoc test, a-b p<0.0001). All experiments were performed at least twice in independent biological repeats with similar results.

**Supplementary Fig. 8:**
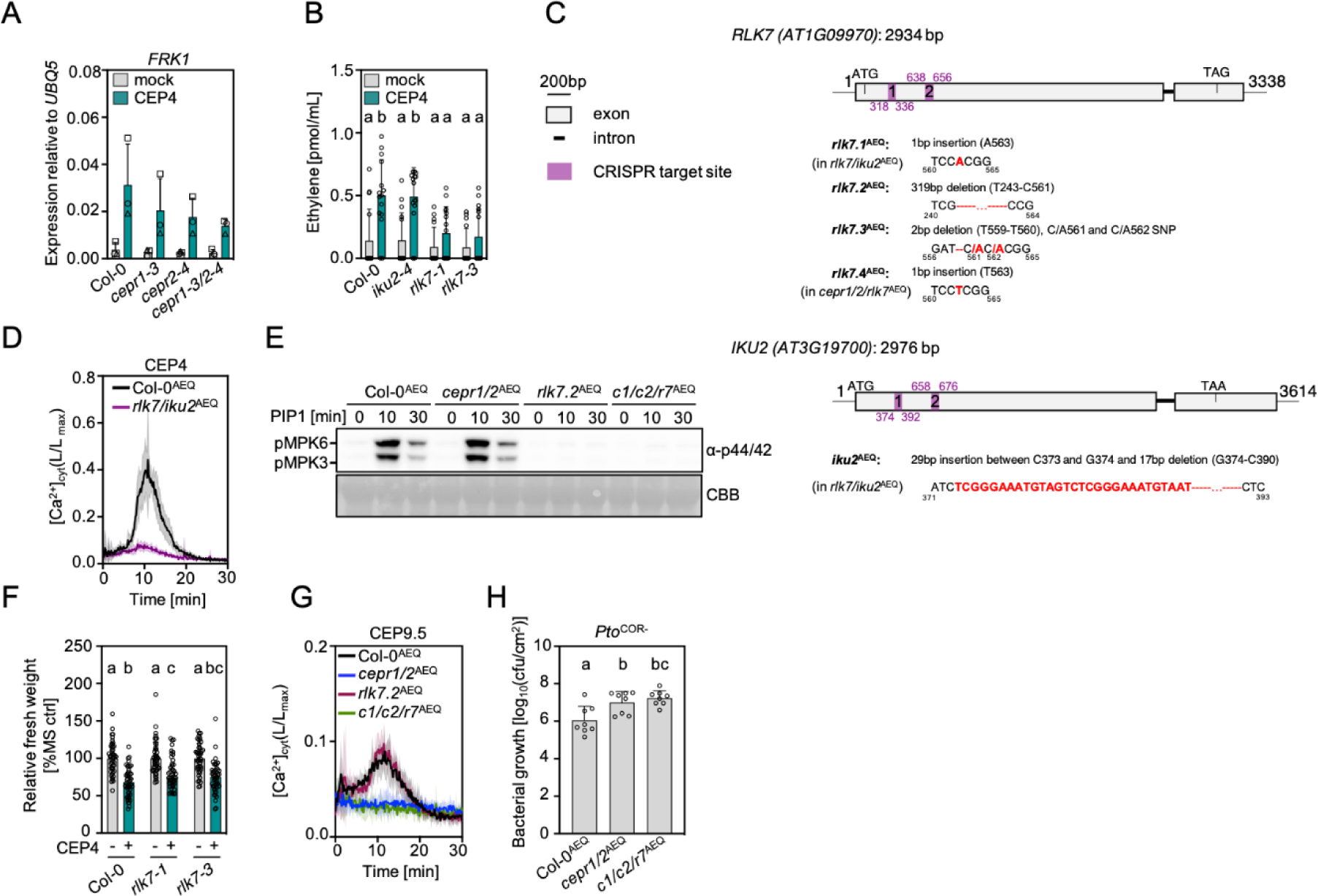
Identification of RLK7 as an additional CEP4 receptor. **A)** RT-qPCR of *FRK1* in seedlings upon mock (ddH_2_O) or CEP4 (1 µM) treatment for 4 h. Housekeeping gene *UBQ5*. Different symbols represent three independent biological repeats ± SD. **B)** CEP4-triggered ethylene production is reduced in *rlk7* mutants. Ethylene concentration in leaf discs upon mock (ddH_2_O) or CEP4 (1 μM) treatment for 3.5 h; n = 15 pooled from three experiments ± SD (one-way ANOVA, Tukey post hoc, a-b p<0.01). **C)** Schematic diagram of the *RLK7* and *IKU2* genomic sequence, structure, and the CRISPR-Cas9-mediated mutation pattern detected by DNA sequencing. The locus number and the length of the coding sequence (CDS) are indicated above the scheme for each gene. There are four *rlk7* alleles generated in Col-0^AEQ^ background: *rlk7.1*^AEQ^ in *rlk7/iku2*^AEQ^ double mutant, two single *rlk7*^AEQ^ mutants: *rlk7.2*^AEQ^ and *rlk7.3*^AEQ^, and *rlk7.4* ^AEQ^ in CRISPR *cepr1/2/rlk7^AEQ^*. CRISPR *iku2* (in *rlk7/iku2*^AEQ^) has a 29 bp insertion, followed by a 17 bp deletion. Mutations in both genes lead to a frameshift mutation and an early stop codon. The specific location and type of mutations for each gene are indicated in the schematics describing the mutants. The two CRISPR target sites are indicated in purple. **D)** [Ca^2+^]_cyt_ kinetics in seedlings upon CEP4 treatment (1 μM); n = 6, ± SD. **E)** MAPK activation upon CEP4 (1 μM) treatment. Western blots were probed with α-p44/42. CBB = Coomassie brilliant blue. **F)** Relative fresh weight of five-day-old seedlings treated with CEP4 (1 μM) for seven days; n = 48 pooled from four experiments ± SD (one-way ANOVA, Tukey post-hoc test, a-b/c/bc p<0.0001, b-c p<0.05). **G)** [Ca^2+^]_cyt_ kinetics in seedlings upon CEP9.5 (10 μM) treatment; n = 6, ± SD. **H)** cfu of *Pto*^COR-^ (3 dpi) upon spray infection; n = 8 pooled from two experiments ± SD (one-way ANOVA, Tukey post-hoc test; a-b, p<0.0120; a-bc p=0.0019). All experiments were performed at least twice in independent biological repeats with similar results.

**Supplementary Fig. 9:**
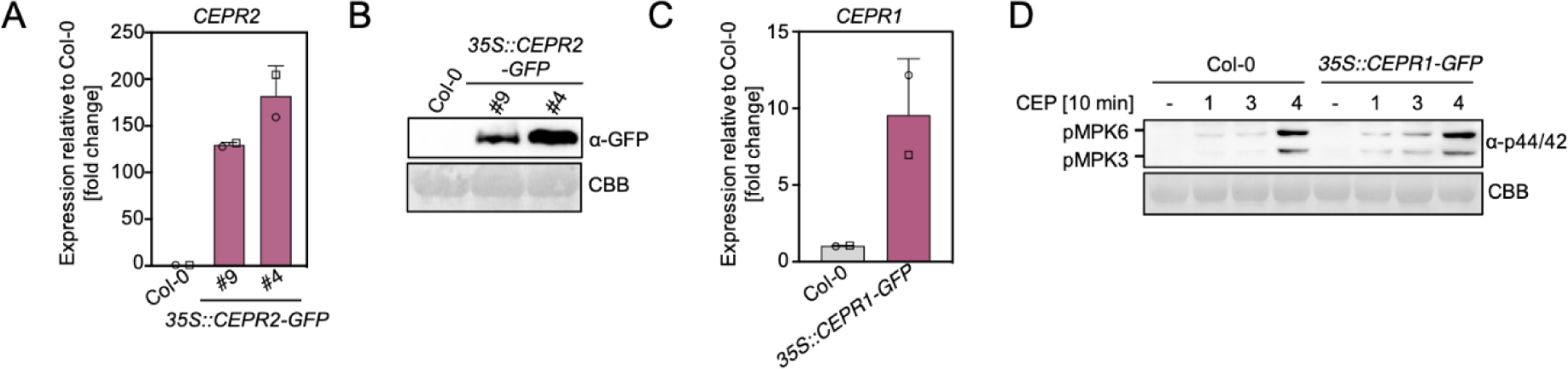
Characterization of *35S::CEPR1-GFP* and *35S::CEPR2-GFP* overexpression lines. **A)** *CEPR2* transcript levels in two independent *CEPR2-GFP* overexpression lines. *CEPR2* expression was normalized to *UBQ5* and is shown as fold induction compared to Col-0. **B)** CEPR2-GFP protein levels in wild-type Col-0 and *CEPR2* overexpression lines. Western blots were probed with α-GFP. CBB = Coomassie brilliant blue. **C)** *CEPR1* transcript levels in the *CEPR1-GFP* overexpression line. *CEPR1* expression was normalized to *UBQ5* and is shown as fold induction compared to Col-0. **D)** MAPK activation in Col-0 and *35S::CEPR1-GFP* line upon CEP1 (1), CEP3 (3) and CEP4 (4) 1 μM treatment. Western blots were probed with α-p44/42. CBB = Coomassie brilliant blue. Different symbols in **A** and **C** represent two independent experiments. All experiments were performed at least twice in independent biological repeats with similar results.

**Supplementary Fig. 10:**
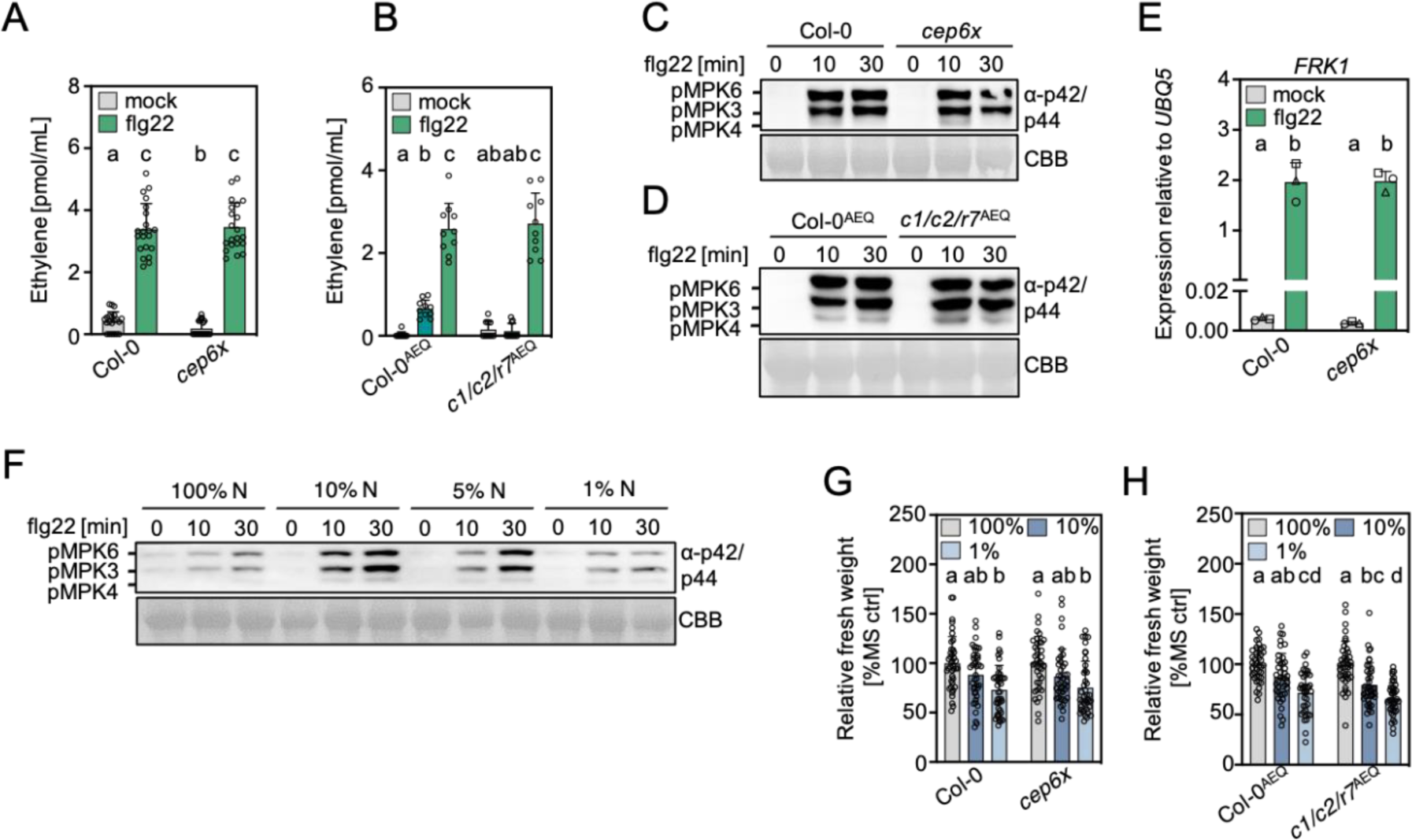
Further characterization of CRISPR *cep* and CRISPR *cepr1/2/rlk7^AEQ^* mutants. **A)** Flg22-triggered ethylene production is not impaired in *cep6x*. Ethylene concentration in leaf discs upon mock (ddH_2_O) or flg22 (500 nM) treatment for 3.5 h; n = 21 pooled from four experiments ± SD (one-way ANOVA, Tukey post hoc, a-b p<0.05; a/b-c p<0.0001). **B)** CEP4-triggered ethylene production is abolished in *cepr1/2/rlk7^AEQ^*. Ethylene concentration in leaf discs upon mock (ddH_2_O), CEP4 (1 μM) or flg22 (500 nM) treatment for 3.5 h; n = 10 pooled from two experiments ± SD (one-way ANOVA, Tukey post hoc, a-b p=0.0166; a/b/ab-c p<0.0001). *cep6x* **(C)** and *cepr1/2/rlk7*^AEQ^ **(D)** are not affected in flg22-triggered MAPK activation under normal conditions. MAPK activation upon flg22 (100 nM) treatment. Western blots were probed with α-p44/42. CBB = Coomassie brilliant blue. **E)** *cep6x* is not affected in flg22-triggered expression of *FRK1* in seedlings. RT-qPCR of *FRK1* upon flg22 (100 nM) treatment for 4 h normalized to *UBQ5*. Different symbols represent different experiments; n = 3 ± SD (one-way ANOVA, Tukey post hoc, a-b p<0.0001). **F)** N availability modulates flg22-triggered MAPK activation. MAPK activation upon flg22 (100 nM) treatment after 24 h transfer of seedlings to 100% N, 10% N, 5% N and 1% N medium. Western blots were probed with α-p44/42. CBB = Coomassie brilliant blue. **G)** The *cep6x* mutant shows wild-type growth response to different medium N concentration. Relative fresh weight of five-day-old seedlings grown in 100% N, 10% N and 1% N medium for seven days; n = 40 pooled from five independent experiments ± SD (one-way ANOVA, Tukey post-hoc test, a-b p<0.001). **H**) The *cepr1/2/rlk7*^AEQ^ shows slightly stronger growth response to different medium N concentrations. Relative fresh weight of five-day-old seedlings grown in 100% N, 10% N and 1% N medium for seven days; n = 39-40 pooled from five independent experiments ± SD (one-way ANOVA, Tukey post-hoc test, a-cd p<0.0001, a-bc p=0.0002, a-d p<0.0001, ab-d, p<0.0001, bc-d p=0.0303).

**Supplementary Table 1:** Peptides used in this study. Hyp (in red) indicates hydroxylated proline residues.

| Peptide name | Sequence |
| --- | --- |
| CEP1 | DFR<sup>Hyp</sup>TNPGNS<sup>Hyp</sup>GVGH |
| CEP2 | DFA<sup>Hyp</sup>TNPGDS<sup>Hyp</sup>GIRH |
| CEP3 | TFR<sup>Hyp</sup>TEPGHS<sup>Hyp</sup>GIGH |
| CEP4<sup>scr</sup> | TGQ<sup>Hyp</sup>DHQR<sup>Hyp</sup>FAHIGGS |
| CEP4<sup>15mer</sup> | AFR<sup>Hyp</sup>THQG<sup>Hyp</sup>SQGIGH |
| CEP4<sup>16mer</sup> | DAFR<sup>Hyp</sup>THQG<sup>Hyp</sup>SQGIGH |
| CEP6.2 | DFEPTT<sup>Hyp</sup>GHS<sup>Hyp</sup>GVGH |
| CEP9.1 | DFV<sup>Hyp</sup>TS<sup>Hyp</sup>GNS<sup>Hyp</sup>GVGH |
| CEP9.2 | DFA<sup>Hyp</sup>TS<sup>Hyp</sup>GHS<sup>Hyp</sup>GVGH |
| CEP9.3 | DFA<sup>Hyp</sup>TS<sup>Hyp</sup>GNS<sup>Hyp</sup>GIGH |
| CEP9.4 | DFA<sup>Hyp</sup>TT<sup>Hyp</sup>GNS<sup>Hyp</sup>GMGH |
| CEP9.5 | DFKPTT<sup>Hyp</sup>GHS<sup>Hyp</sup>GVGH |
| PIP1 | RLASG<sup>Hyp</sup>SPRGRGH |
| flg22 | Ac-QRLSTGSRINSAKDDAAGLQIA |

**Supplementary Table 2:**
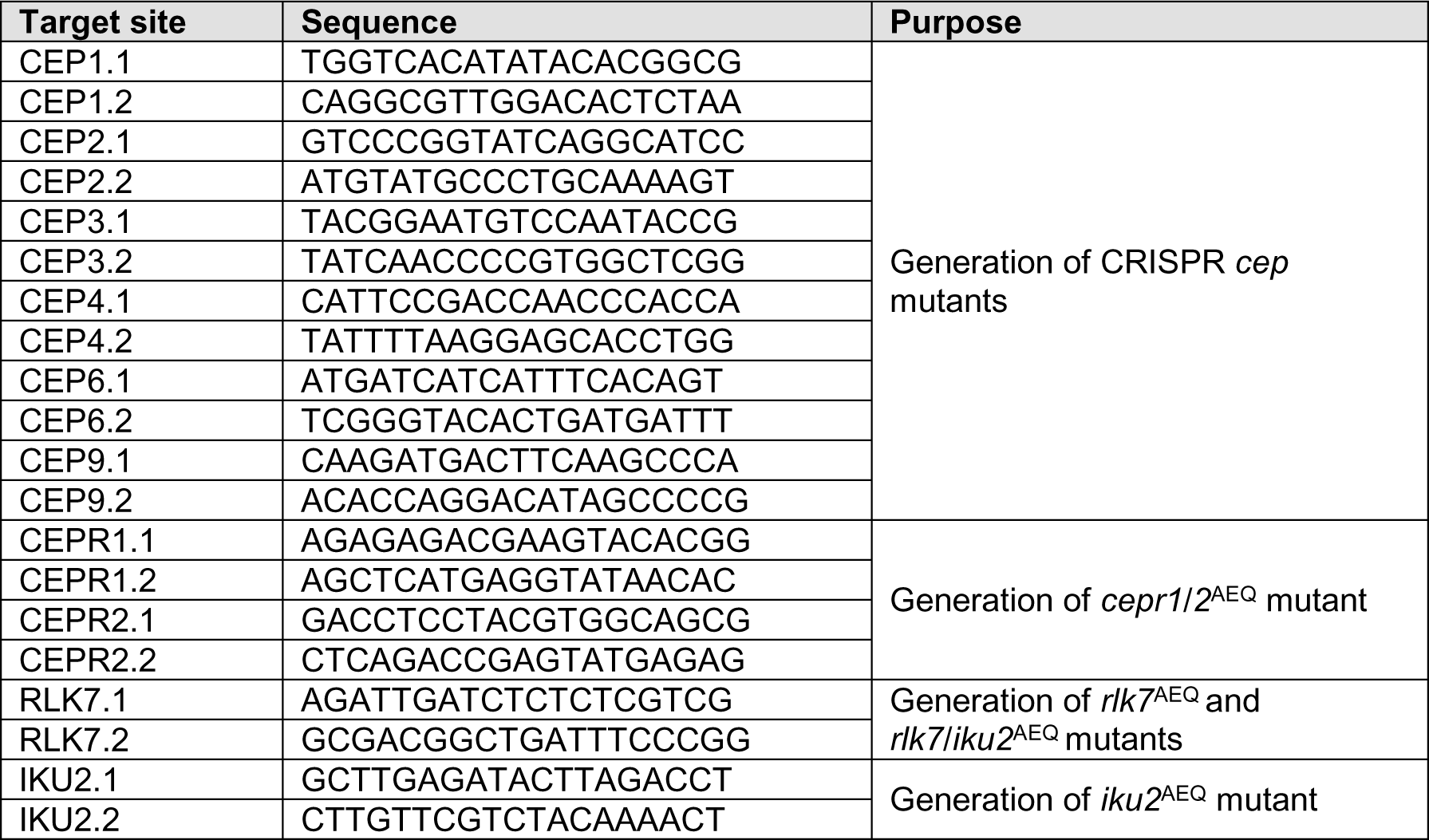
CRISPR-Cas9 target sites.

**Supplementary Table 3:**
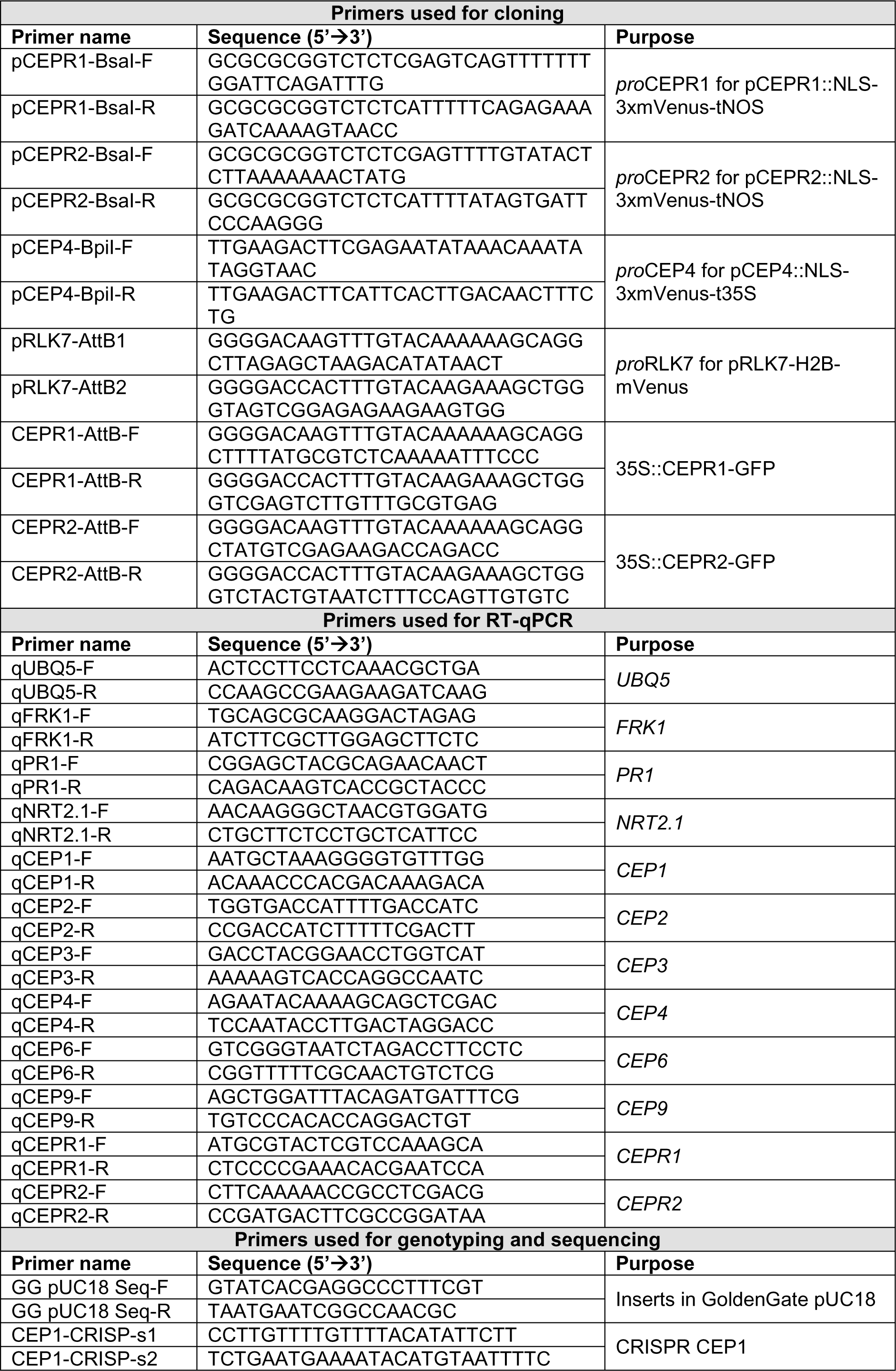
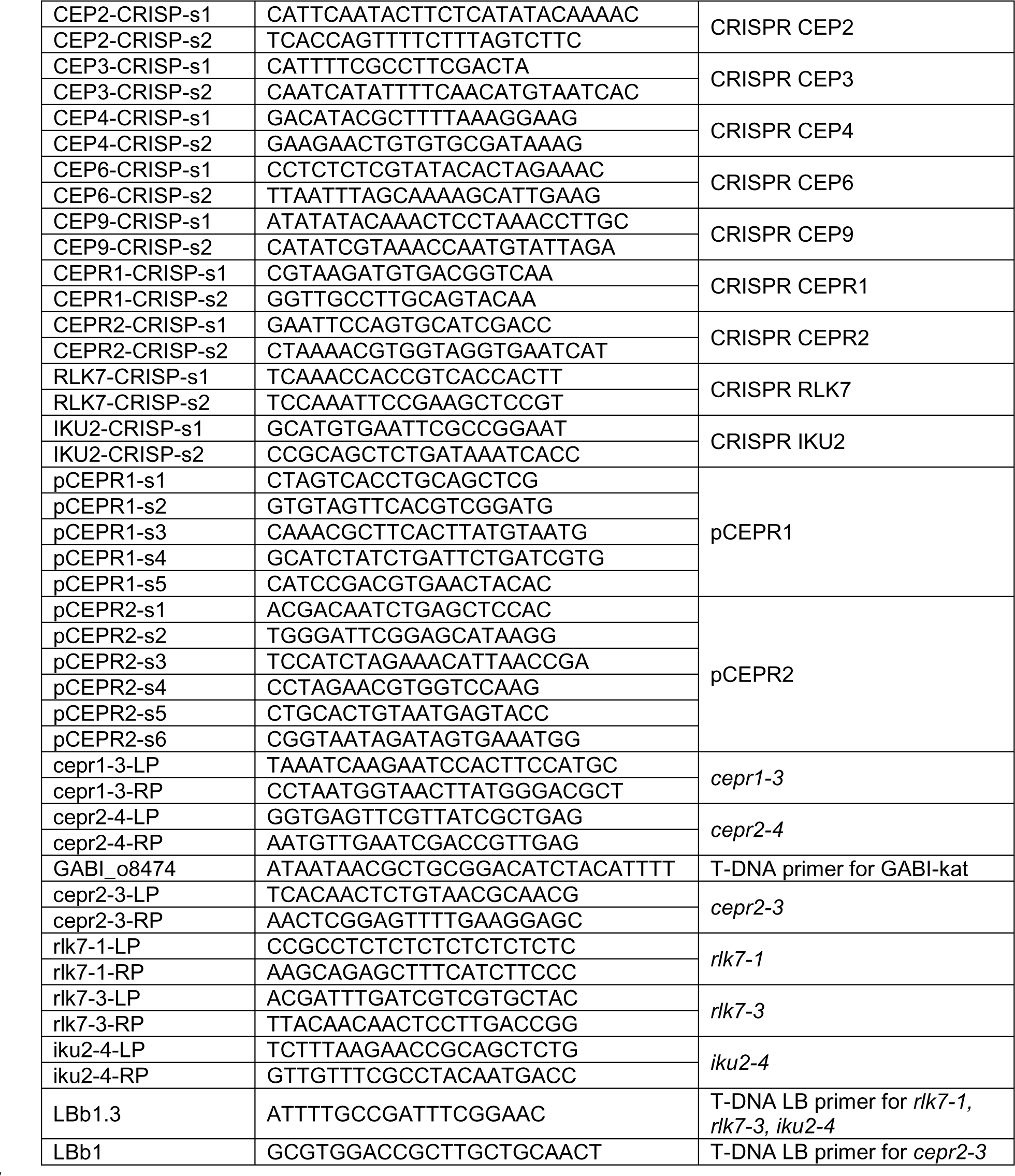
Primers used in this study.

## References

1. L. Gómez-Gómez, G. Felix, T. Boller, A single locus determines sensitivity to bacterial flagellin in Arabidopsis thaliana. Plant J. 18, 277–84 (1999).

2. C. Zipfel, S. Robatzek, L. Navarro, E. J. Oakeley, J. D. G. Jones, G. Felix, T. Boller, Bacterial disease resistance in Arabidopsis through flagellin perception. Nature. 428, 764–767 (2004).

3. A. Heese, D. R. Hann, S. Gimenez-Ibanez, A. M. E. Jones, K. He, J. Li, J. I. Schroeder, S. C. Peck, J. P. Rathjen, The receptor-like kinase SERK3/BAK1 is a central regulator of innate immunity in plants. Proc. Natl. Acad. Sci. USA. 104, 12217–22 (2007).

4. D. Chinchilla, C. Zipfel, S. Robatzek, B. Kemmerling, T. Nürnberger, J. D. G. Jones, G. Felix, T. Boller, A flagellin-induced complex of the receptor FLS2 and BAK1 initiates plant defence. Nature. 448, 497–500 (2007).

5. V. Olsson, L. Joos, S. Zhu, K. Gevaert, M. A. Butenko, I. De Smet, Look closely, the beautiful may be small: precursor-derived peptides in plants. Annu. Rev. Plant Biol. 70, 1–1.34 (2019).

6. A. A. Gust, R. Pruitt, T. Nürnberger, Sensing danger: Key to activating plant immunity. Trends Plant Sci. 22, 779–791 (2017).

7. J. Rzemieniewski, M. Stegmann, Regulation of pattern-triggered immunity and growth by phytocytokines. Curr. Opin. Plant Biol. 68, 102230 (2022).

8. M. Stegmann, P. Zecua-Ramirez, C. Ludwig, H.-S. Lee, B. Peterson, Z. L. Nimchuk, Y. Belkhadir, R. Hückelhoven, RGI-GOLVEN signalling promotes cell surface immune receptor abundance to regulate plant immunity. EMBO Rep. 23, e53281 (2022).

9. M. Stegmann, J. Monaghan, E. Smakowska-Luzan, H. Rovenich, A. Lehner, N. Holton, Y. Belkhadir, C. Zipfel, The receptor kinase FER is a RALF-regulated scaffold controlling plant immune signaling. Science. 355, 287–289 (2017).

10. Y. Xiao, M. Stegmann, Z. Han, T. A. DeFalco, K. Parys, L. Xu, Y. Belkhadir, C. Zipfel, J. Chai, Mechanisms of RALF peptide perception by a heterotypic receptor complex. Nature. 572, 270– 274 (2019).

11. J. Gronnier, C. M. Franck, M. Stegmann, T. DeFalco, A. Abarca, M. Von Arx, K. Duenser, W. Lin, Z. Yang, J. Kleine-Vehn, C. Ringli, C. Zipfel, Regulation of immune receptor kinases plasma membrane nanoscale landscape by a plant peptide hormone and its receptors. Elife. 11, e74162 (2022).

12. R. Whitford, A. Fernandez, R. Tejos, A. C. Pérez, J. Kleine-Vehn, S. Vanneste, A. Drozdzecki, J. Leitner, L. Abas, M. Aerts, K. Hoogewijs, P. Baster, R. De Groodt, Y. C. Lin, V. Storme, Y. Van de Peer, T. Beeckman, A. Madder, B. Devreese, C. Luschnig, J. Friml, P. Hilson, GOLVEN secretory peptides regulate auxin carrier turnover during plant gravitropic responses. Dev. Cell. 22, 678–685 (2012).

13. M. R. Blackburn, M. Haruta, D. S. Moura, Twenty Years of Progress in Physiological and Biochemical Investigation of RALF peptides. Plant Physiol. 182, 1657–1666 (2020).

14. S. Zhu, Q. Fu, F. Xu, H. Zheng, F. Yu, New paradigms in cell adaptation: decades of discoveries on the CrRLK1L receptor kinase signalling network. New Phytol. 232, 1168–1183 (2021).

15. C. Liu, L. Shen, Y. Xiao, D. Vyshedsky, C. Peng, X. Sun, Z. Liu, L. Cheng, H. Zhang, Z. Han, J. Chai, H. Wu, A. Y. Cheung, C. Li, Pollen PCP-B peptides unlock a stigma peptide – receptor kinase gating mechanism for pollination. Science. 372, 171–175 (2021).

16. S. Zhong, L. Li, Z. Wang, Z. Ge, Q. Li, A. Bleckmann, J. Wang, Z. Song, Y. Shi, T. Liu, L. Li, H. Zhou, Y. Wang, L. Zhang, H.-M. Wu, L. Lai, H. Gu, J. Dong, A. Y. Cheung, T. Dresselhaus, L.-J. Qu, RALF peptide signaling controls the polytubey block in Arabidopsis. Science. 375, 290– 296 (2022).

17. Z. Lan, Z. Song, Z. Wang, L. Li, Y. Liu, S. Zhi, R. Wang, J. Wang, Q. Li, Antagonistic RALF peptides control an intergeneric hybridization barrier on Brassicaceae stigmas. Cell. 186, 1–15 (2023).

18. J. Huang, L. Yang, L. Yang, X. Wu, X. Cui, L. Zhang, J. Hui, Y. Zhao, H. Yang, S. Liu, Q. Xu, M. Pang, X. Guo, Y. Cao, Y. Chen, X. Ren, J. Lv, J. Yu, J. Ding, G. Xu, N. Wang, X. Wei, Q. Lin, Y. Yuan, X. Zhang, C. Ma, C. Dai, P. Wang, Y. Wang, F. Cheng, W. Zeng, R. Palanivelu, X. Zhang, A. Y. Cheung, Q. Duan, Stigma receptors control intraspecies and interspecies barriers in Brassicaceae. Nature. 614, 303–308 (2022).

19. K. Gully, S. Pelletier, M.-C. C. Guillou, M. Ferrand, S. Aligon, I. Pokotylo, A. Perrin, E. Vergne, M. Fagard, E. Ruelland, P. Grappin, E. Bucher, J.-P. P. Renou, S. Aubourg, The SCOOP12 peptide regulates defense response and root elongation in Arabidopsis thaliana. J. Exp. Bot. 70, 1349–1365 (2019).

20. J. Rhodes, A.-O. Roman, M. Bjornson, B. Brandt, P. Derbyshire, M. Wyler, M. Schmid, F. L. H. Menke, J. Santiago, C. Zipfel, Perception of a conserved family of plant signalling peptides by the receptor kinase HSL3. Elife. 11, e74687 (2022).

21. Z. Liu, S. Hou, O. Rodrigues, P. Wang, D. Luo, K. Nomura, C. Yin, H. Wang, W. Zhang, K. Zhu-salzman, Phytocytokine signalling reopens stomata in plant immunity and water loss. Nature. 605, 332–339 (2022).

22. R. Tabata, K. Sumida, T. Yoshii, K. Ohyama, H. Shinohara, Y. Matsubayashi, Perception of root-derived peptides by shoot LRR-RKs mediates systemic N-demand signaling. Science. 346, 343– 346 (2014).

23. F. Takahashi, T. Suzuki, Y. Osakabe, S. Betsuyaku, Y. Kondo, N. Dohmae, H. Fukuda, K. Yamaguchi-Shinozaki, K. Shinozaki, A small peptide modulates stomatal control via abscisic acid in long-distance signalling. Nature. 556, 235–238 (2018).

24. H. Zhou, F. Xiao, Y. Zheng, G. Liu, Y. Zhuang, Z. Wang, Y. Zhang, J. He, C. Fu, H. Lin, PAMP-INDUCED SECRETED PEPTIDE 3 modulates salt tolerance through RECEPTOR-LIKE KINASE 7 in plants. Plant Cell. 34, 927–944 (2021).

25. K. Ohyama, M. Ogawa, Y. Matsubayashi, Identification of a biologically active, small, secreted peptide in Arabidopsis by in silico gene screening, followed by LC-MS-based structure analysis. Plant J. 55, 152–160 (2008).

26. M. Taleski, N. Imin, M. A. Djordjevic, CEP peptide hormones: Key players in orchestrating nitrogen-demand signalling, root nodulation, and lateral root development. J. Exp. Bot. 69, 1829–1836 (2018).

27. C. Delay, K. Chapman, M. Taleski, Y. Wang, S. Tyagi, Y. Xiong, N. Imin, M. A. Djordjevic, CEP3 levels affect starvation-related growth responses of the primary root. J. Exp. Bot. 70, 4763–4774 (2019).

28. K. Chapman, M. Taleski, H. A. Ogilvie, N. Imin, M. A. Djordjevic, CEP-CEPR1 signalling inhibits the sucrose-dependent enhancement of lateral root growth. J. Exp. Bot. 70, 3955–3967 (2019).

29. S. Roy, L. M. Müller, A rulebook for peptide control of legume – microbe endosymbioses. Trends Plant Sci. 27, 870–889 (2022).

30. K. Chapman, M. Taleski, M. Frank, M. A. Djordjevic, CEP and cytokinin hormone signaling intersect to promote shallow lateral root angles. J. Exp. Bot. in press, erad353 (2023).

31. M. Taleski, K. Chapman, O. Novák, T. Schmülling, M. Frank, M. A. Djordjevic, CEP peptide and cytokinin pathways converge on CEPD glutaredoxins to inhibit root growth. Nat. Commun. 13, 1683 (2023).

32. B. Li, S. Jiang, X. Yu, C. Cheng, S. Chen, Y. Cheng, J. S. Yuan, D. Jiang, P. He, L. Shan, Phosphorylation of Trihelix Transcriptional Repressor ASR3 by MAP KINASE4 Negatively Regulates Arabidopsis Immunity. Plant Cell. 27, 839–856 (2015).

33. I. Roberts, S. Smith, B. De Rybel, J. Van Den Broeke, W. Smet, S. De Cokere, M. Mispelaere, I. De Smet, T. Beeckman, The CEP family in land plants: Evolutionary analyses, expression studies, and role in Arabidopsis shoot development. J. Exp. Bot. 64, 5371–5381 (2013).

34. M. Melotto, W. Underwood, J. Koczan, K. Nomura, S. Y. He, Plant Stomata Function in Innate Immunity against Bacterial Invasion. Cell. 126, 969–980 (2006).

35. V. Nekrasov, J. Li, M. Batoux, M. Roux, Z.-H. Chu, S. Lacombe, A. Rougon, P. Bittel, M. Kiss-Papp, D. Chinchilla, H. P. van Esse, L. Jorda, B. Schwessinger, V. Nicaise, B. P. H. J. Thomma, A. Molina, J. D. G. Jones, C. Zipfel, Control of the pattern-recognition receptor EFR by an ER protein complex in plant immunity. EMBO J. 28, 3428–38 (2009).

36. M. R. Knight, A. K. Campbellt, S. M. Smith, A. J. Trewavas, Transgenic plant aequorin reports the effects of touch and cold-shock and elicitors on cytoplasmic calcium. Nature. 314, 524–526 (1991).

37. L. Huang, Y. Yuan, C. Lewis, L. Dandurand, I. Zasada, L. Huang, Y. Yuan, C. Lewis, J. Kud, J. C. Kuhl, A. Caplan, L. Dandurand, NILR1 perceives a nematode ascaroside triggering immune signaling and resistance. Curr. Biol. 33, 1–6 (2023).

38. L. Poncini, I. Wyrsch, V. D. Tendon, T. Vorley, T. Boller, N. Geldner, J. P. Métraux, S. Lehmann, In roots of Arabidopsis thaliana, the damage-associated molecular pattern AtPep1 is a stronger elicitor of immune signalling than flg22 or the chitin heptamer. PLoS One. 12, e0185808 (2017).

39. C. Delay, N. Imin, M. A. Djordjevic, CEP genes regulate root and shoot development in response to environmental cues and are specific to seed plants. J. Exp. Bot. 64, 5383–5394 (2013).

40. C. Furumizu, A. K. Krabberød, M. Hammerstad, R. M. Alling, M. Wildhagen, S. Sawa, R. B. Aalen, The sequenced genomes of nonflowering land plants reveal the innovative evolutionary history of peptide signaling. Plant Cell. 33, 2915–2934 (2021).

41. M. Taleski, K. Chapman, N. Imin, M. A. Djordjevic, M. Groszmann, The peptide hormone receptor CEPR1 functions in the reproductive tissue to control seed size and yield. Plant Physiol. 182, 620–636 (2020).

42. K. Chapman, A. Ivanovici, M. Taleski, C. J. Sturrock, J. L. P. Ng, N. A. Mohd-Radzman, F. Frugier, M. J. Bennett, U. Mathesius, M. A. Djordjevic, CEP receptor signalling controls root system architecture in Arabidopsis and Medicago. New Phytol. 226, 1809–1821 (2020).

43. J. Rhodes, H. Yang, S. Moussu, F. Boutrot, J. Santiago, C. Zipfel, Perception of a divergent family of phytocytokines by the Arabidopsis receptor kinase MIK2. Nat. Commun. 12, 5494 (2021).

44. P. Qian, W. Song, T. Yokoo, A. Minobe, G. Wang, T. Ishida, S. Sawa, J. Chai, T. Kakimoto, The CLE9/10 secretory peptide regulates stomatal and vascular development through distinct receptors. Nat. Plants. 4, 1071–1081 (2018).

45. S. Hou, D. Liu, S. Huang, D. Luo, Z. Liu, Q. Xiang, P. Wang, R. Mu, Z. Han, S. Chen, J. Chai, L. Shan, P. He, The Arabidopsis MIK2 receptor elicits immunity by sensing a conserved signature from phytocytokines and microbes. Nat. Commun. 12, 5494 (2021).

46. F. Zhu, J. Deng, H. Chen, P. Liu, L. Zheng, Q. Ye, R. Li, M. Brault, J. Wen, F. Frugier, J. Dong, T. Wang, A CEP Peptide Receptor-like Kinase Regulates Auxin Biosynthesis and Ethylene Signaling to Coordinate Root Growth and Symbiotic Nodulation in Medicago truncatula. Plant Cell. 32, 2855–2877 (2020).

47. D. Mackey, B. F. Holt, A. Wiig, J. L. Dangl, RIN4 interacts with Pseudomonas syringae type III effector molecules and is required for RPM1-mediated resistance in Arabidopsis. Cell. 108, 743– 54 (2002).

48. H. H. Breitenbach, M. Wenig, F. Wittek, L. Jordá, A. M. Maldonado-Alconada, H. Sarioglu, T. Colby, C. Knappe, M. Bichlmeier, E. Pabst, D. Mackey, J. E. Parker, A. Corina Vlot, Contrasting roles of the apoplastic aspartyl protease APOPLASTIC, ENHANCED DISEASE SUSCEPTIBILITY1-DEPENDENT1 and LEGUME LECTIN-LIKE PROTEIN1 in Arabidopsis systemic acquired resistance. Plant Physiol. 165, 791–809 (2014).

49. M. Luo, E. S. Dennis, F. Berger, W. J. Peacock, A. Chaudhury, MINISEED3 (MINI3), a WRKY family gene, and HAIKU2 (IKU2), a leucine-rich repeat (LRR) KINASE gene, are regulators of seed size in Arabidopsis. Proc. Natl Acad. Sci. USA. 102, 17531–17536 (2005).

50. K. Toyokura, T. Goh, H. Shinohara, A. Shinoda, Y. Kondo, Y. Okamoto, T. Uehara, K. Fujimoto, Y. Okushima, Y. Ikeyama, K. Nakajima, T. Mimura, M. Tasaka, Y. Matsubayashi, H. Fukaki, Lateral inhibition by a peptide hormone-receptor cascade during Arabidopsis lateral root founder cell formation. Dev. Cell. 48, 64–75.e5 (2019).

51. D. Pitorre, C. Llauro, E. Jobet, J. Guilleminot, J. P. Brizard, M. Delseny, E. Lasserre, RLK7, a leucine-rich repeat receptor-like kinase, is required for proper germination speed and tolerance to oxidative stress in Arabidopsis thaliana. Planta. 232, 1339–1353 (2010).

52. S. Hou, X. Wang, D. Chen, X. Yang, M. Wang, D. Turrà, A. Di Pietro, W. Zhang, The secreted peptide PIP1 amplifies immunity through Receptor-Like Kinase 7. PLoS Pathog. 10, e1004331 (2014).

53. S. Hou, H. Shen, H. Shao, PAMP-induced peptide 1 cooperates with salicylic acid to regulate stomatal immunity in Arabidopsis thaliana. Plant Signal. Behav. 14, 1666657 (2019).

54. Y. Wu, Q. Xun, Y. Guo, J. Zhang, K. Cheng, T. Shi, K. He, S. Hou, X. Gou, J. Li, Genome-Wide Expression Pattern Analyses of the Arabidopsis Leucine-Rich Repeat Receptor-Like Kinases. Mol. Plant. 9, 289–300 (2016).

55. Y. Ohkubo, M. Tanaka, R. Tabata, M. Ogawa-Ohnishi, Y. Matsubayashi, Shoot-to-root mobile polypeptides involved in systemic regulation of nitrogen acquisition. Nat. Plants. 3, 17029 (2017).

56. R. Ota, Y. Ohkubo, Y. Matsubayashi, M. Ogawa-ohnishi, Shoot-to-root mobile CEPD-like 2 integrates shoot nitrogen status to systemically regulate nitrate uptake in Arabidopsis. Nat. Commun. 11, 641 (2020).

57. E. Hoffland, M. J. Jeger, M. L. van Beusichem, Effect of nitrogen supply rate on disease resistance in tomato depends on the pathogen. Plant Soil. 218, 239–247 (2000).

58. M. Fagard, A. Launay, G. Clément, J. Courtial, A. Dellagi, M. Farjad, A. Krapp, M. C. Soulié, C. Masclaux-Daubresse, Nitrogen metabolism meets phytopathology. J. Exp. Bot. 65, 5643–5656 (2014).

59. C. Verly, A. C. R. Djoman, M. Rigault, F. Giraud, L. Rajjou, M. E. Saint-Macary, A. Dellagi, Plant defense stimulator mediated defense activation is affected by nitrate fertilization and developmental stage in Arabidopsis thaliana. Front. Plant Sci. 11, 583 (2020).

60. D. Zhuo, M. Okamoto, J. J. Vidmar, A. D. M. Glass, Regulation of a putative high-affinity nitrate transporter (Nrt2;1At) in roots of Arabidopsis thaliana. Plant J. 17, 563–568 (1999).

61. T. Kiba, A. B. Feria-Bourrellier, F. Lafouge, L. Lezhneva, S. Boutet-Mercey, M. Orsel, V. Bréhaut, A. Miller, F. Daniel-Vedele, H. Sakakibara, A. Krapp, S. Figure, S. Figure, The Arabidopsis nitrate transporter NRT2.4 plays a double role in roots and shoots of nitrogen-starved plants. Plant Cell. 24, 245–258 (2012).

62. Z. Luo, J. Wang, F. Li, Y. Lu, Z. Fang, M. Fu, K. S. K. S. Mysore, J. Wen, J. Gong, J. D. Murray, F. Xie, The small peptide CEP1 and the NIN-like protein NLP1 regulate NRT2.1 to mediate root nodule formation across nitrate concentrations. Plant Cell. 35, 776–794 (2023).

63. A. Roman, P. Jimenez-sandoval, S. Augustin, C. Broyart, L. A. Hothorn, J. Santiago, HSL1 and BAM1/2 impact epidermal cell development by sensing distinct signaling peptides. Nat. Commun. 13, 876 (2022).

64. J. Dindas, T. A. DeFalco, G. Yu, L. Zhang, P. David, M. Bjornson, M.-C. Thibaud, V. Custódio, G. Castrillo, L. Nussaume, A. P. Macho, C. Zipfel, Direct inhibition of phosphate transport by immune signaling in Arabidopsis. Curr. Biol. 32, 488–495.e5 (2022).

65. S. Roy, M. Griffiths, I. Torres-Jerez, B. Sanchez, E. Antonelli, D. Jain, N. Krom, S. Zhang, L. M. York, W. R. Scheible, M. Udvardi, Application of Synthetic Peptide CEP1 Increases Nutrient Uptake Rates Along Plant Roots. Front. Plant Sci. 12, 793145 (2022).

66. E. A. Vidal, J. M. Alvarez, V. Araus, E. Riveras, M. D. Brooks, G. Krouk, S. Ruffel, L. Lejay, N. M. Crawford, G. M. Coruzzi, R. A. Gutiérrez, Nitrate in 2020: Thirty years from transport to signaling networks. Plant Cell. 32, 2094–2119 (2020).

67. C. H. Ho, S. H. Lin, H. C. Hu, Y. F. Tsay, CHL1 Functions as a Nitrate Sensor in Plants. Cell. 138, 1184–1194 (2009).

68. K. Liu, M. Liu, Z. Lin, Z. Wang, B. Chen, C. Liu, A. Guo, M. Konishi, S. Yanagisawa, G. Wagner, J. Sheen, NIN-like protein 7 transcription factor is a plant nitrate sensor. Science. 377, 1419– 1425 (2022).

69. A. Jacquot, V. Chaput, A. Mauries, Z. Li, P. Tillard, C. Fizames, P. Bonillo, F. Bellegarde, E. Laugier, V. Santoni, S. Hem, A. Martin, A. Gojon, W. Schulze, L. Lejay, NRT2.1 C-terminus phosphorylation prevents root high affinity nitrate uptake activity in Arabidopsis thaliana. New Phytol. 228, 1038–1054 (2020).

70. Y. Ohkubo, K. Kuwata, Y. Matsubayashi, A type 2C protein phosphatase activates high-affinity nitrate uptake by dephosphorylating NRT2.1. Nat. Plants. 7, 310–316 (2021).

71. X. Zou, M. Y. Liu, W. H. Wu, Y. Wang, Phosphorylation at Ser28 stabilizes the Arabidopsis nitrate transporter NRT2.1 in response to nitrate limitation. J. Integr. Plant Biol. 62, 865–876 (2019).

72. W. Wang, B. Hu, A. Li, C. Chu, NRT1.1s in plants: Functions beyond nitrate transport. J. Exp. Bot. 71, 4373–4379 (2020).

73. L. Zhang, Z. Yu, Y. Xu, M. Yu, Y. Ren, S. Zhang, G. Yang, J. Huang, K. Yan, C. Zheng, C. Wu, Regulation of the stability and ABA import activity of NRT1.2/NPF4.6 by CEPR2-mediated phosphorylation in Arabidopsis. Mol. Plant. 14, 633–646 (2021).

74. C. Laffont, A. Ivanovici, P. Gautrat, M. Brault, M. A. Djordjevic, F. Frugier, The NIN transcription factor coordinates CEP and CLE signaling peptides that regulate nodulation antagonistically. Nat. Commun. 11, 3167 (2020).

75. A. Ivanovici, C. Laffont, E. Larrainzar, N. Patel, C. S. Winning, H. C. Lee, N. Imin, F. Frugier, M. A. Djordjevic, The Medicago SymCEP7 hormone increases nodule number via shoots without compromising lateral root number. Plant Physiol. 191, 2012–2026 (2023).

76. M. Taleski, M. Jin, K. Chapman, K. Taylor, C. Winning, M. Frank, N. Imin, M. A. Djordjevic, CEP hormones at the nexus of nutrient acquisition and allocation, root development, and plant-microbe interactions. J. Exp. Bot. in press, erad444 (2023).

77. B. Castel, L. Tomlinson, F. Locci, Y. Yang, J. D. G. Jones, Optimization of T-DNA architecture for Cas9-mediated mutagenesis in Arabidopsis. PLoS One. 14, e0204778 (2019).

78. M. Somssich, A. Bleckmann, R. Simon, Shared and distinct functions of the pseudokinase CORYNE (CRN) in shoot and root stem cell maintenance of Arabidopsis. J. Exp. Bot. 67, 4901– 4915 (2016).

79. I. Dimitrov, F. E. Tax, Lateral root growth in Arabidopsis is controlled by short and long distance signaling through the LRR RLKs XIP1/CEPR1 and CEPR2. Plant Signal. Behav. 13, e1489667 (2018).

80. A. Fatihi, A. M. Zbierzak, P. Dörmann, Alterations in seed development gene expression affect size and oil content of Arabidopsis seeds. Plant Physiol. 163, 973–985 (2013).

81. M. Roux, B. Schwessinger, C. Albrecht, D. Chinchilla, A. Jones, N. Holton, F. G. Malinovsky, M. Tör, S. de Vries, C. Zipfel, The Arabidopsis leucine-rich repeat receptor-like kinases BAK1/SERK3 and BKK1/SERK4 are required for innate immunity to hemibiotrophic and biotrophic pathogens. Plant Cell. 23, 2440–55 (2011).

82. F. Zhou, A. Emonet, V. Dénervaud Tendon, P. Marhavy, D. Wu, T. Lahaye, N. Geldner, Co-incidence of damage and microbial patterns controls localized immune responses in roots. Cell. 180, 440–453 (2020).

83. C. W. Melnyk, Grafting with arabidopsis thaliana. Methods Mol. Biol. 1497, 9–18 (2017).

84. M. Wenig, A. Ghirardo, J. H. Sales, E. S. Pabst, H. H. Breitenbach, F. Antritter, B. Weber, B. Lange, M. Lenk, R. K. Cameron, J. P. Schnitzler, A. C. Vlot, Systemic acquired resistance networks amplify airborne defense cues. Nat. Commun. 10, 3813 (2019).

85. M. Futatsumori-Sugai, K. Tsumoto, Signal peptide design for improving recombinant protein secretion in the baculovirus expression vector system. Biochem. Biophys. Res. Commun. 391, 931–935 (2010).

86. Y. Hashimoto, S. Zhang, S. Zhang, Y. R. Chen, G. W. Blissard, Correction: BTI-Tnao38, a new cell line derived from Trichoplusia ni, is permissive for AcMNPV infection and produces high levels of recombinant proteins. BMC Biotechnol. 12 (2012), doi:10.1186/1472-6750-12-12.

